# Rifampicin tolerance and growth fitness among isoniazid-resistant clinical *Mycobacterium tuberculosis* isolates: an in-vitro longitudinal study

**DOI:** 10.1101/2023.11.22.568240

**Authors:** Srinivasan Vijay, Nguyen Le Hoai Bao, Dao Nguyen Vinh, Le Thanh Hoang Nhat, Do Dang Anh Thu, Nguyen Le Quang, Le Pham Tien Trieu, Hoang Ngoc Nhung, Vu Thi Ngoc Ha, Phan Vuong Khac Thai, Dang Thi Minh Ha, Nguyen Huu Lan, Maxine Caws, Guy E. Thwaites, Babak Javid, Nguyen Thuy Thuong Thuong

## Abstract

Antibiotic tolerance in *Mycobacterium tuberculosis* leads to less effective bacterial killing, poor treatment responses and resistant emergence. Therefore, we investigated the rifampicin tolerance of *M. tuberculosis* isolates, with or without pre-existing isoniazid-resistance. We determined the *in-vitro* rifampicin survival fraction by minimum duration of killing assay in isoniazid susceptible (IS, n=119) and resistant (IR, n=84) *M. tuberculosis* isolates. Then we correlated the rifampicin tolerance with bacterial growth, rifampicin minimum inhibitory concentrations (MICs) and isoniazid-resistant mutations. The longitudinal IR isolates collected from patients were analyzed for changes in rifampicin tolerance and associated emergence of genetic variants. The median duration of rifampicin exposure reducing the *M. tuberculosis* surviving fraction by 90% (minimum duration of killing-MDK90) increased from 1.23 (95%CI 1.11; 1.37) and 1.31 (95%CI 1.14; 1.48) to 2.55 (95%CI 2.04; 2.97) and 1.98 (95%CI 1.69; 2.56) days, for IS and IR respectively, during 15 to 60 days of incubation. This indicated the presence of fast and slow growing tolerant sub-populations. A range of 6 log_10_-fold survival fraction enabled classification of tolerance as low, medium or high and revealed IR association with increased tolerance with faster growth (OR=2.68 for low vs. medium, OR=4.42 for low vs. high, *P*-trend=0.0003). The high tolerance in IR isolates was specific to those collected during rifampicin treatment in patients and associated with bacterial genetic microvariants. Furthermore, the high rifampicin tolerant IR isolates have survival potential similar to multi-drug resistant isolates. These findings suggest that IR tuberculosis needs to be evaluated for high rifampicin tolerance to improve treatment regimen and prevent the risk of MDR-TB emergence.

## Introduction

*Mycobacterium tuberculosis* causes around 10 million cases of tuberculosis (TB) each year and 1.5 million deaths^1^. Challenges to successful TB treatment include bacterial evolution and diversification under host stresses and antibiotics, leading to differential antibiotic susceptibility even among genetically-susceptible *M. tuberculosis* isolates^2^. Based on killing dynamics, the differential susceptibility can be classified into two phenomena, 1) antibiotic tolerance observed as reduced rate of killing of the entire bacterial population^3^, and 2) antibiotic persistence observed as reduced rate of killing of sub-populations compared to more susceptible bacteria^4,5^. Clinically susceptible isolates exposed to host stresses and antibiotic selection can exhibit increased antibiotic tolerance and persistence^6–8^, as seen by the emergence of mutations increasing tolerance or persistence among clinical *M. tuberculosis* isolates^9–12^. Recent studies have also implicated the antibiotic tolerance in clinical isolates as a risk factor for hard-to treat infections and tolerance can also contribute to the emergence of resistance^13^ and relapse^14^.

Emergence of rifampicin tolerance or persistence, a key drug in TB treatment is a major concern considering the emergence of multi-drug resistant (MDR, resistant to at least isoniazid and rifampicin) tuberculosis^15^. Several mechanisms lead to rifampicin tolerance, heteroresistance or persistence^16^. These include efflux pump overexpression^17^, mistranslation^18^, overexpression of rifampicin target *rpoB*^19^, cell size heterogeneity^20–22^ and the redox heterogeneity in bacteria^23^. Rifampicin treatment can also result in differentially detectable sub-populations of *M. tuberculosis*, which can grow only in liquid medium as compared to solid medium^24^. Therefore, in determining risk of post-treatment relapse, it is important to consider, alongside tolerance range, the degree of growth heterogeneity within tolerant subpopulations.

Apart from rifampicin susceptibility variation, another concern in standard TB treatment is the emergence of isoniazid resistance (IR). There is globally around 10% prevalence of IR among clinical *M. tuberculosis* isolates^25^. IR is difficult to rapidly diagnose during drug susceptibility testing, and is associated with worse treatment outcomes compared to isoniazid-susceptible (IS) *M. tuberculosis* isolates^25^. Importantly, IR is also associated with subsequent emergence of rifampicin resistance leading to MDR TB^26^.

Despite its potential importance for the TB treatment, the distribution of rifampicin tolerance among clinical *M. tuberculosis* isolates is unknown, and routine clinical microbiology diagnosis does not include any assays for tolerance. The growth fitness of rifampicin tolerant subpopulations, and the association of pre-existing IR with rifampicin tolerance is completely unknown.

To address this knowledge gap, we developed a most-probable number (MPN) based minimum duration of killing (MDK) assay to determine the rifampicin tolerance among clinical *M. tuberculosis* isolates in a medium-throughput manner^27^. In the current study, we investigated the rifampicin tolerance in a large set of IS (n=119) and IR (n=84) clinical *M. tuberculosis* isolates and its correlation with bacterial growth rate, rifampicin MICs, IR-mutations and the rifampicin treatment selection in patients.

## Results

### Study design

We investigated rifampicin tolerance and its association with isoniazid susceptibility among 242 clinical *M. tuberculosis* isolates. We treated susceptible isolates with rifampicin (2µg/mL), a concentration several times higher than their MICs (supplementary table 1) and also close to the serum rifampicin concentration observed in patient during oral dose^28^, and at 0, 2 and 5 days determined fractional survival following 15, 30 and 60 days of culture (figure 1A). Higher survival fractions indicate higher rifampicin tolerance, and differences in survival fraction determined between 15 and 60 days of incubation indicated greater growth heterogeneity in rifampicin tolerant sub-populations (figure 1B). 23 of the isolates grew poorly in the absence of antibiotic, and a further 10 had low initial MPN, making accurate determination of survival fractions difficult (figure 1 A), and these 33 isolates were removed from further analysis. Of the remaining 209 isolates, 119 IS, 84 IR and 6 resistant to both rifampicin and isoniazid, MDR. The MDR isolates were controls and comparators as isolates with a known high degree of rifampicin tolerance^27^.

**Figure 1.**
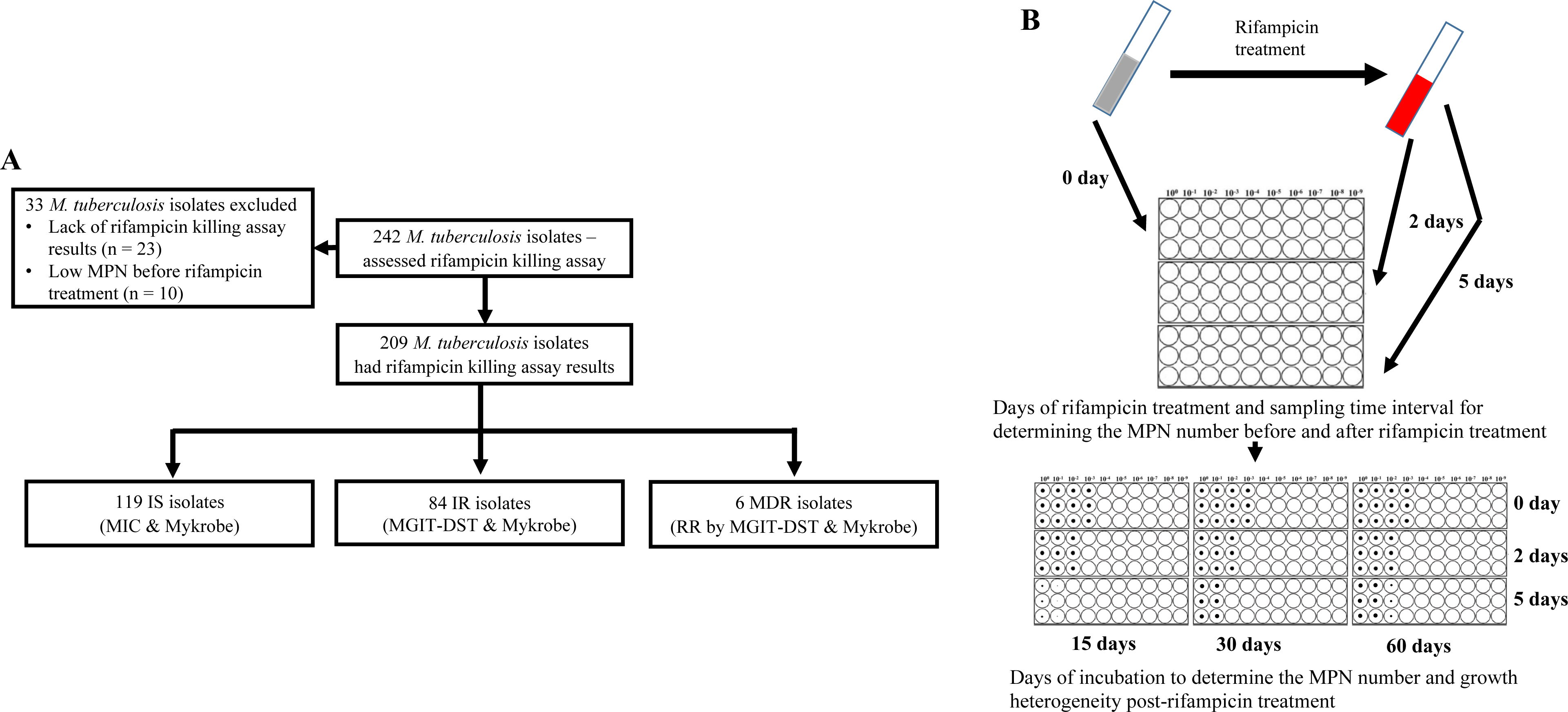
Study design. (A) Study design. IS – Isoniazid susceptible, IR – Isoniazid resistant, RR – Rifampicin resistant. (B) Most-probable number based rifampicin killing assay and survival fraction determination.

### Distribution of Rifampicin tolerance in IS and IR isolates

We analyzed the rifampicin survival fraction and the kill curve for IS and IR *M. tuberculosis* isolates, at 0, 2 and 5 days of rifampicin treatment followed by 15 and 60 days of incubation (figure 2). We did not further analyse 30 days incubation result, as it was similar to 60 days incubation (supplementary figure 1). Following 5 days of rifampicin treatment, the average survival fraction reduced by 90-99% of the starting bacterial population (figure 2). We calculated the time required for 90% survival fraction reduction (MDK_90_) for each isolate by determine the different length of X-axis (Days post rifampicin treatment) corresponding to 90% decline in survival fraction in Y-axis (figure 2, supplementary figure 2). Of note, the average time required for 90% survival fraction reduction (MDK_90_) was 1.23 (95%CI (Confidence interval) 1.11; 1.37) and 1.31 (95%CI 1.14; 1.48) days for IS and IR respectively when survivors were incubated for 15 days, but rose to 2.55 (95%CI 2.04; 2.97) and 1.98 (95%CI 1.69; 2.56) days for 60 days for IS and IR isolates respectively (figure 2). This shift in the MDK_90_ indicated the presence of growth heterogeneity within the tolerant subpopulation – with both fast and slow-growing bacteria within tolerant subpopulations. For most of the isolates MDK_90_ time could be calculated but other parameters of tolerance and persistence such as MDK_99_ (at 15 day=81% (170/209), 60 day=41% (86/209)) and MDK_99.99_ (at 15 day=11% (22/209), 60 day=8% (17/209)) could be calculated for only a fraction of 209 isolates and the rest were beyond the assay limits (supplementary figure 2). Intriguingly, we observed a significant difference in rifampicin tolerance between IS and IR isolates at 5 days of treatment– but only in the 15 days post recovery. The difference had disappeared by 60 days (figure 2). Therefore, we decided to consider survival fractions with 15 and 60 days recovery for further analysis, the earliest and latest time points for determining the fast- and slowly-growing rifampicin tolerant subpopulations.

**Figure 2.**
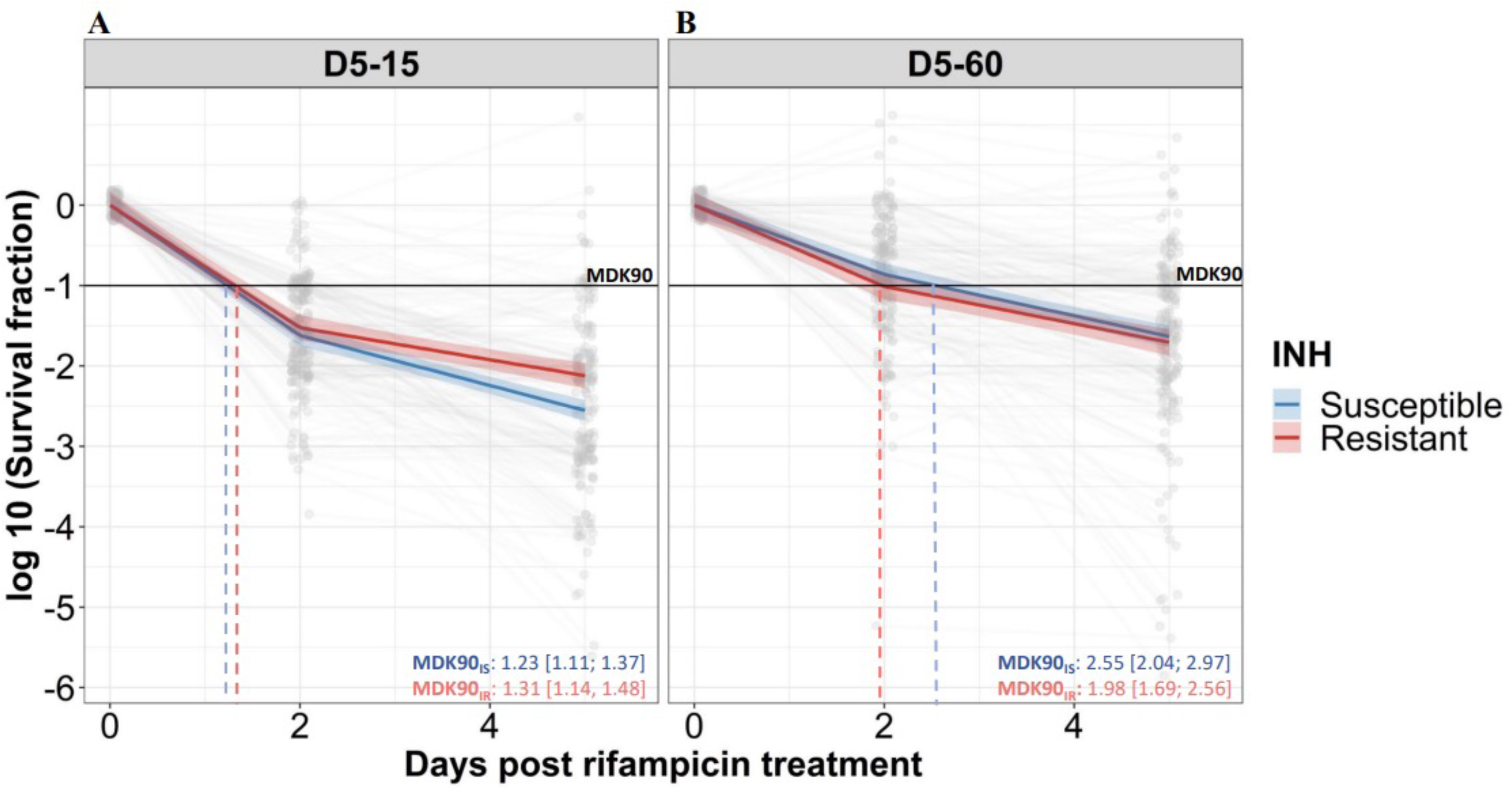
Rifampicin survival curve in isoniazid susceptible and resistant clinical *M. tuberculosis* isolates. **(A, B)** The bacterial kill curve as measured by log_10_ survival fraction from data collected at 0, 2 and 5 days of rifampicin treatment followed by incubation for 15 and 60 days respectively. Data from individual isolates are shown as the grey dots connected by lines. Estimated mean with 95% credible interval (bold coloured line and colour shaded area respectively) of isoniazid susceptible (IS – blue, n = 119, 117 for 15 and 60 days of incubation respectively) and resistant (IR – red, n = 84, 80 for 15 and 60 days of incubation respectively) clinical *M. tuberculosis* isolates based on linear mixed effect model implemented in a Bayesian framework. One log_10_ fold or 90% reduction in survival fraction is indicated (MDK90, black horizontal line) and the mean time duration required for 90% reduction in survival (MDK90, minimum duration of killing time) at 15 and 60 days of incubation is indicated by vertical dashed lines with respective colours for IS and IR isolates.

### Isoniazid resistance is associated with fast-growing rifampicin tolerant subpopulations

To further group rifampicin tolerance level, and correlate it with growth fitness and isoniazid susceptibility, we compared the distribution of survival fraction at 15 and 60 days recovery following 2 and 5 days of rifampicin treatment in IS (n=119) and IR (n=84) isolates (figure 3A, supplementary figure 3). There was no significant difference in rifampicin tolerance between IS and IR isolates at 2 days of treatment (supplementary figure 3). At 5 days of rifampicin treatment and both early (15 days) and late (60 days) recovery time points, IS and IR isolates showed a broad distribution of fractional survival–spanning 1 million-times difference in rifampicin susceptibility (figure 3 A). At the 15 days recovery period, IR was significantly associated with higher survival to rifampicin treatment as compared to IS isolates (P=0.017, figure 3A), whereas at 60 days, fractional survival increased in both groups with no difference according to isoniazid susceptibility (figure 3A). These results suggest that the difference between IS and IR rifampicin tolerant subpopulations is within their fast-growing tolerant bacilli only.

**Figure 3.**
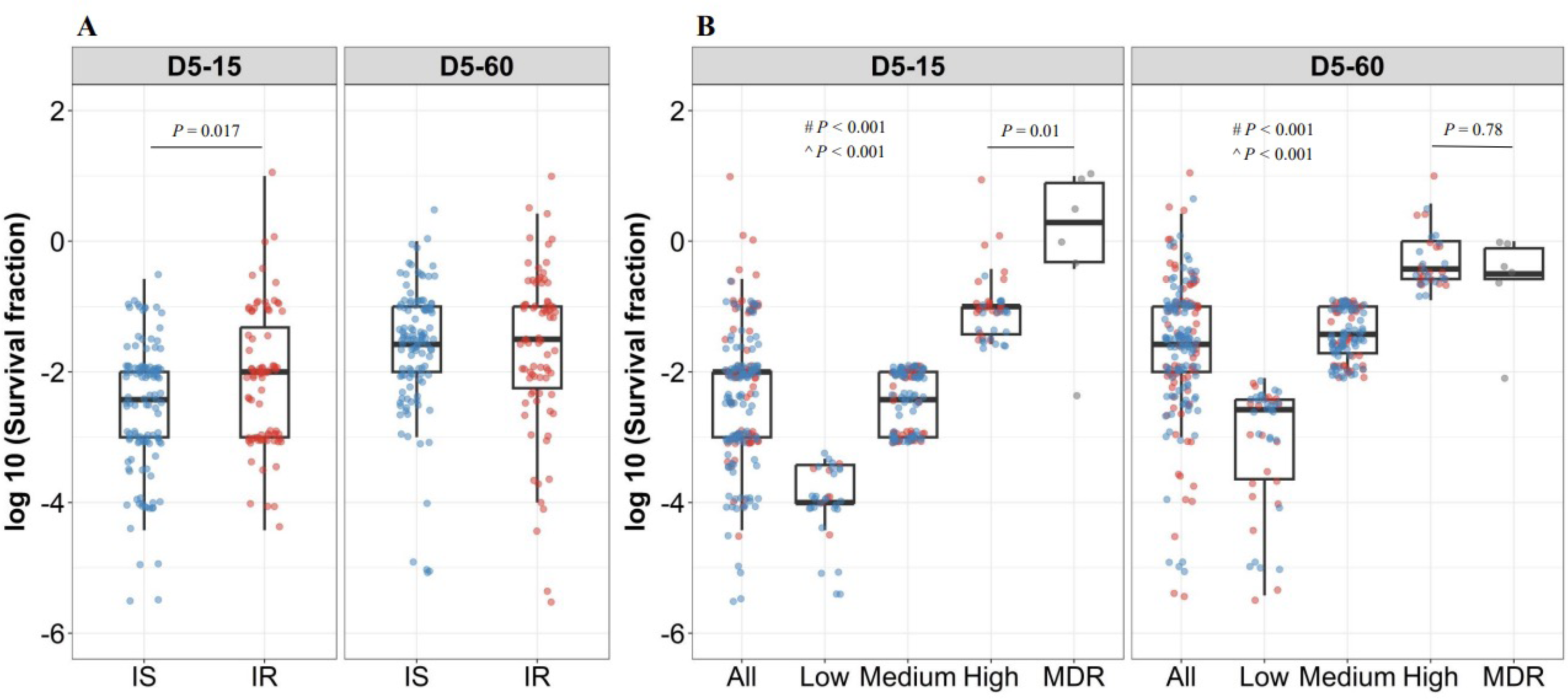
Rifampicin survival fraction distribution in isoniazid susceptible and resistant clinical *M. tuberculosis* isolates. (A) Log_10_ rifampicin survival fraction distribution, with median and IQR (inter quartile range), of individual isoniazid susceptible (IS, blue dots, n = 119, 117 for D5-15, and D5-60 respectively), and resistant (IR, red dots, n = 84, 80 for D5-15, D5-60 respectively) isolates for 5 days of rifampicin treatment as determined at 15 and 60 days of incubation (D5-15, D5-60 respectively). (B) Rifampicin tolerance distribution in both IS (blue dots) and IR (red dots) isolates combined together (All) was used to group them as low (< 25th percentile, n = 33, 47 for D5-15, and D5-60 respectively), medium (from 25th to 75th percentile, n = 124, 115 for D5-15, and D5-60 respectively) and high (above 75th percentile, n = 46, 35 for D5-15, and D5-60 respectively) level of rifampicin tolerance and compare it with rifampicin tolerance of MDR clinical *M. tuberculosis* isolates (grey dots, n = 6), after 5 days of rifampicin treatment and determined at 15 and 60 days of incubation (D5-15, D5-60 respectively). Statistical comparisons between Low, Medium, and High or MDR were made by using Wilcoxon rank-sum test. # - p-value for comparing the Low and High tolerance groups, ^ - p-value for comparing the medium and High tolerance groups.

To further refine distribution of rifampicin tolerance between isolates, first we combined the rifampicin survival fraction distribution of both IS and IR isolates, then the fractional rifampicin survival was parsed as low, medium or high as defined by falling within the 25^th^, 75^th^ and 100^th^ percentiles of survival fractions following rifampicin treatment and either 15 or 60 days recovery (figure 3B). As expected, there was substantially lower tolerance to rifampicin in low and medium groups compared with MDR isolates. Surprisingly, tolerance to rifampicin between non-rifampicin resistant “high” tolerance strains and MDR strains was not significantly different (P=0.78, figure 3B), and these high tolerant strains were characterized in both IS and IR isolates. This suggests that within the IR, high tolerant subgroup, antibiotic susceptibility (to both rifampicin and isoniazid) may be similar to *bona fide* MDR strains.

Analyzing rifampicin tolerance subgroups between IS and IR strains, at the early, 15 day recovery time-point, the majority (79%, 26/33) of “low” rifampicin tolerant strains were isoniazid susceptible. By contrast, IR isolates were over-represented in the “medium” and “high” tolerant subgroups (OR of 2.7 and 4.4 respectively–table 1). These associations disappeared with longer (60 day) recovery post antibiotic treatment, confirming that IR isolates harbored fast-growing, high-level rifampicin-tolerant bacilli compared with IS isolates (table 1).

**Table 1.** Association of rifampicin tolerance level with isoniazid susceptibility.

| Incubation time | Rifampicin tolerance level | Isoniazid Susceptible (N = 119) | Isoniazid Resistant (N = 84) | P | OR (95%CI) | P trend |
| --- | --- | --- | --- | --- | --- | --- |
| D5-15 | Low tolerance (N, %) | 26 (79, 26/33) | 7 (21, 7/33) |  |  | 0.0038 |
|  | Medium tolerance (N, %) | 72 (58, 72/124) | 52 (42, 52/124) | 0.029 | 2.68 (1.08-6.65) |  |
|  | High tolerance (N, %) | 21 (46, 21/46) | 25 (54, 25/46) | 0.003 | 4.42 (1.60-12.22) |  |
| D5-60 | Low tolerance (N, %) | 26 (55, 26/47) | 21 (45, 21/47) |  |  | 0.67 |
|  | Medium tolerance (N, %) | 74 (64, 74/115) | 41 (36, 41/115) | 0.28 | 0.69 (0.34-1.37) |  |
|  | High tolerance (N, %) | 17 (49, 17/35) | 18 (51, 18/35) | 0.55 | 1.31 (0.55-3.15) |  |

N = number of isolates. (% as percentage, N/total number (IS + IR))
P = P-value determined using Chi-square test.
P trend = P- value determined using Cochran-Armitage test.
OR = odds ratio.
95%CI = 95% confidence interval.

### Association between rifampicin tolerance and relative growth in absence of antibiotics, rifampicin MICs, isoniazid resistant mutations of *M. tuberculosis* isolates

Clinical isolates of *M. tuberculosis* exhibit a large degree of lag time and growth heterogeneity^29^, as well as differences in rifampicin MICs or isoniazid-resistant mutations. Prior studies showed that slow growth rate and non-replicating persistence were correlated^30^, therefore we wished to investigate the association between growth rates in the absence of antibiotic treatment, rifampicin MIC distribution, isoniazid-resistant mutations and rifampicin tolerance distribution in *M. tuberculosis* isolates.

For correlating relative growth in absence of antibiotics, we removed 19 outliers which deviated from normal distribution (supplementary figure 4 with 19 outliers), Intriguingly, slower growth before rifampicin treatment did not have significant correlation with higher growth fitness in rifampicin survival fraction at 15 days incubation in IS isolates (figure 4A regression coefficient −0.21, 95%CI [-0.44, 0.007], *P*=0.058). By contrast, correlation of slower growth with lower growth fitness in the long recovery period was observed in both IS and IR isolates (figure 4B, regression coefficient for IS=0.38 [0.15, 0.61], *P*=0.0014, and IR=0.38 [0.12, 0.64], *P*=0.0041). Comparing IS and IR isolates, IR isolates had slower growth in the absence of antibiotics (figure 4C, P<0.0001). Thus, slow growth before rifampicin treatment correlates with reduced growth fitness in certain rifampicin tolerant populations at 60 days incubation.

**Figure 4.**
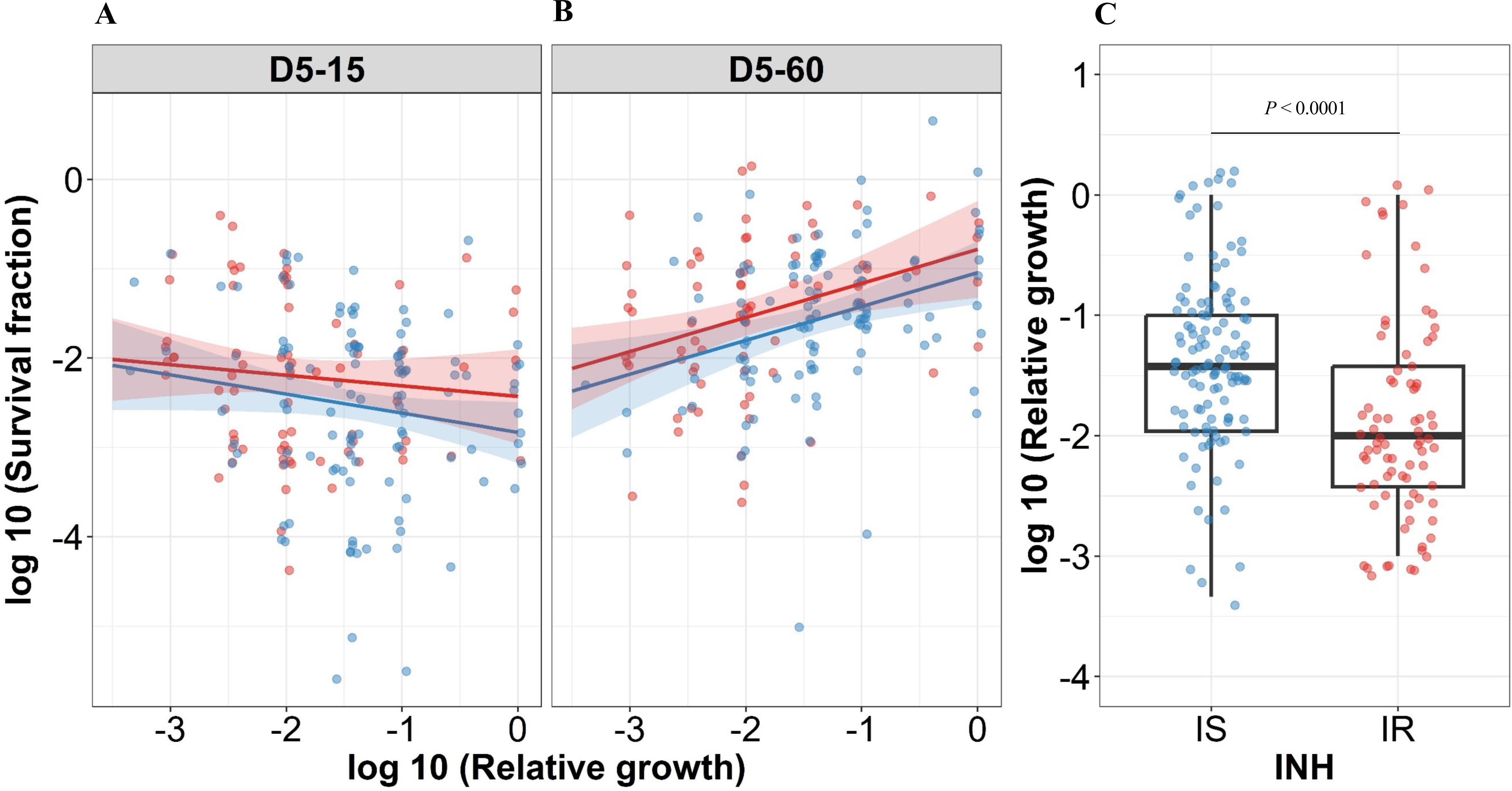
Correlating rifampicin survival fraction with before treatment relative growth of clinical *M. tuberculosis* isolates. Log_10_ survival fraction at 5 days of rifampicin treatment as determined at 15 days (A) and 60 days of incubation (B), for isoniazid susceptible (IS, blue dots) and resistant (IR, red dots) isolates respectively, correlated with the log10 relative growth determined before rifampicin treatment for clinical *M. tuberculosis* isolates. Coefficients of linear regression for (A) IS = - 0.21 [-0.44, 0.007], *P* = 0.058; IR = −0.12 [-0.38, 0.14], *P* = 0.37, and (B) IS = 0.38 [0.15, 0.61], *P* = 0.0014; IR = 0.38 [0.12, 0.64], *P* = 0.0041. (C) Log_10_ distribution of relative growth with median and IQR for IS and IR clinical *M. tuberculosis* isolates before rifampicin treatment. Statistical comparisons between IS and IR were made by using Wilcoxon rank-sum test.

In case of IS isolates, higher rifampicin MICs correlated with lower rifampicin tolerance at long recovery period, 15 (−0.24 [-0.50, 0.022], *P*=0.073) and 60 days incubation (−0.31 [-0.53, −0.083], *P*=0.007, supplementary figure 5A), whereas IR isolates did not show such a negative correlation of rifampicin tolerance with rifampicin MICs (0.14 [-0.14, 0.41], *P*=0.33 and 0.21 [-0.057, 0.48], *P*=0.12, supplementary figure 5A). This latter observation might be due to increased growth fitness of IR rifampicin tolerant populations. In addition, there was no significant difference in rifampicin MICs distribution between IS and IR isolates (supplementary figure 5B).

We next investigated the association between isoniazid-resistant mutations in *M. tuberculosis* isolates and rifampicin tolerance distribution. These isolates had three different isoniazid-resistant mutations, *katG*_S315X (n=71), *inhA*_I21T (n=2) and fabG1_C-15X (n=6) and data not available for 5 isolates (supplementary figure 6). Due to low number of isolates with inhA and fabG1 mutations, it was not possible to identify the difference in rifampicin tolerance distribution between the isolates with different isoniazid-resistant mutations. Nevertheless, we observed wide distribution of rifampicin tolerance in isoniazid-resistant isolates with katG_S315X mutation itself (supplementary figure 6), indicating the role of other genetic or epi-genetic determinants influencing rifampicin tolerance.

### Higher rifampicin tolerance and growth fitness is associated with IR isolates from intensive phase of treatment with rifampicin

The IS isolates were collected only at baseline before treatment, whereas the IR isolates in our study were collected longitudinally from patients at different stages of treatment. Both patients with IS and IR isolates received the standard 8 months treatment regimen according to the Vietnamese National TB Program during the study period^25^, this included initial two months with four antibiotics (streptomycin or ethambutol, with rifampicin, isoniazid and pyrazinamide) followed by 6 months with isoniazid and ethambutol^25^. The antibiotic treatment may select different *M. tuberculosis* genetic microvariants in the patients and lead to differences in rifampicin tolerance between longitudinal isolates. Therefore, we analyzed the rifampicin tolerance distribution in the IR isolates in three sub-groups, before treatment (IR-BL), initial two months of intensive phase of treatment with rifampicin in the regimen (IR-IP), continuous phase and relapse lacking rifampicin and any other antibiotics treatment selection respectively (IR-CP) (Figure 5). This grouping reflects TB-treatment regimen in Vietnam during the study period with rifampicin only in the initial two months of treatment^25^. Interestingly, we observed significantly higher rifampicin tolerance and growth fitness in IR-IP group (*P*=0.0018, Figure 5 as compared to IS, IR-BL and IR-CP groups during 15 days recovery, indicating rifampicin treatment itself as a possible mechanism leading to the selection of *M. tuberculosis* tolerant microvariants in patients^19^.

**Figure 5.**
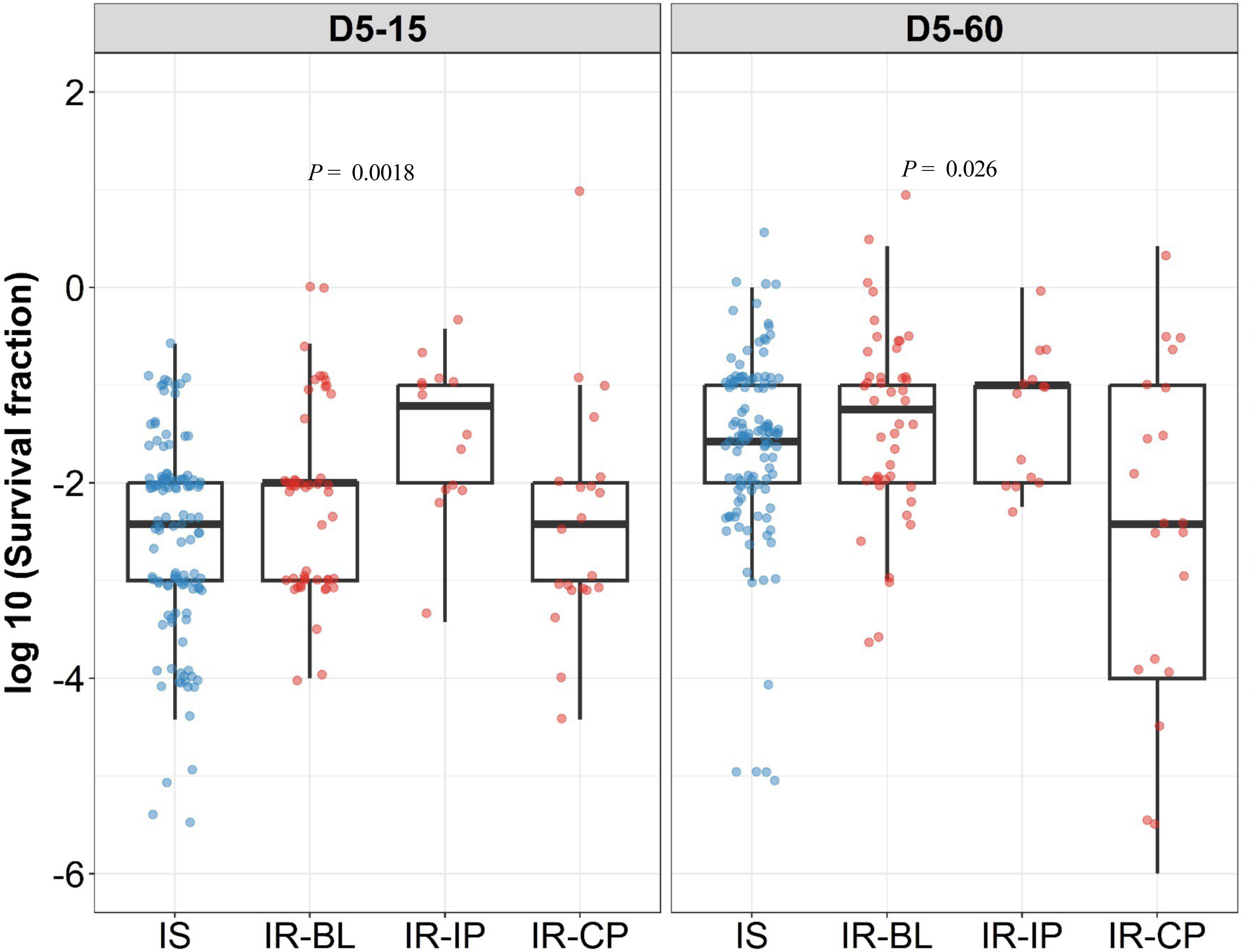
Rifampicin survival fraction distribution in isoniazid susceptible and longitudinal isoniazid resistant clinical *M. tuberculosis* isolates. Log_10_ rifampicin survival fraction distribution, with median and IQR (inter quartile range), of individual isoniazid susceptible (IS, blue dots, n = 119, 117 for D5-15, and D5-60 respectively), and longitudinal isoniazid resistant (IR, red dots, n = 84, 80 for D5-15, D5-60 respectively) isolates for 5 days of rifampicin treatment as determined at 15 and 60 days of incubation (D5-15, D5-60 respectively) grouped based on collection time as baseline (IR-BL, n = 49), intensive phase (IR-IP, n = 14), and continuous phase and relapse (IR-CP, n = 21). Statistical comparisons between groups were made by using Krusal-Walis test.

To verify this finding, we grouped individual patients (n = 18) based on changes in rifampicin tolerance between their initial and subsequent IR isolates collected before treatment (0 month), during treatment (1 to 8 months) and post-treatment (12 to 24 months) (Figure 6). We observed three kinds of changes in rifampicin tolerance between the isolates collected from same patient, 1. decrease (one or more subsequent isolates with lower rifampicin tolerance as compared to the initial isolate), 2. unchanged (initial and subsequent isolates with similar level of rifampicin tolerance) and 3. Increase (one or more subsequent isolates with higher rifampicin tolerance as compared to the initial isolate) for 5 days or rifampicin treatment and 15 and 60 days recovery time (Figure 6) and analyzed the difference in non-synonymous SNPs between the isolates from the same patients associated with differences in rifampicin tolerance (Figure 7, supplementary table 2). The SNPs difference between the longitudinally collected *M. tuberculosis* isolates from same patient were 0-3 except in one case (SNPs=11), indicating de-novo emergence or selection of genetic microvariants within the patient (supplementary table 2). Next, we analyzed the non-synonymous SNPs associated with the changes in rifampicin tolerance both at 15 and 60 days incubation. This included both genetic variants emerging as more than 90% of WGS reads and less-than 90% threshold used as a cut-off for calling SNPs. Several genes Rv0792c, Rv1266c, Rv1696, Rv1758, Rv1997, Rv2043c, Rv2329c, Rv2394, Rv2398c, Rv2400c, Rv2488c, Rv2545, Rv2689c, Rv3138, Rv3680 and Rv3758c previously reported to be associated with persistence, tolerance and survival within host had non-synonymous SNPs associated with changes in rifampicin tolerance (Figure 7, supplementary table 3 with references). This indicates mutations in multiple genes might affect rifampicin tolerance and growth fitness, since there was no one gene or genetic variant in *M. tuberculosis* in multiple patients consistently associated with increased or decreased rifampicin tolerance, or that mutations may be epistatic with the genetic background of the strain.

**Figure 6.**
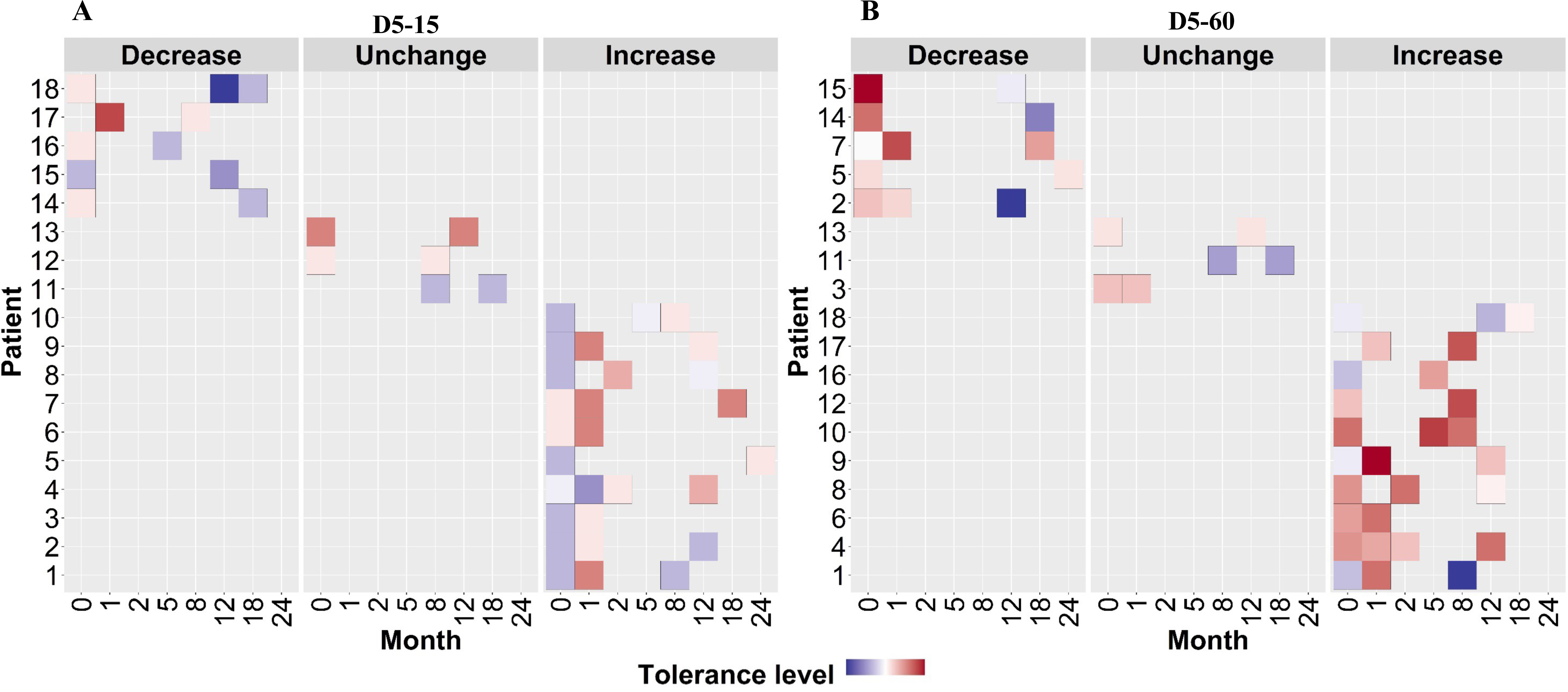
Rifampicin tolerance of longitudinal isoniazid resistant clinical *M. tuberculosis* isolates from individual patients. **(A, B)** Rifampicin tolerance heat map after 5 days of rifampicin treatment as determined at 15 and 60 days of incubation (D5-15, D5-60 respectively), of longitudinal isoniazid resistant clinical *M. tuberculosis* isolates collected from individual patients during different months of treatment and follow-up. Longitudinal isoniazid resistant clinical *M. tuberculosis* isolates from individual patients are grouped based on changes in rifampicin tolerance compared between initial and subsequent months of collection as decrease, un change and increase. Months (0 – 24) represent the different months the isolates were collected from patients during 8 months treatment and 24 months follow-up.

**Figure 7.**
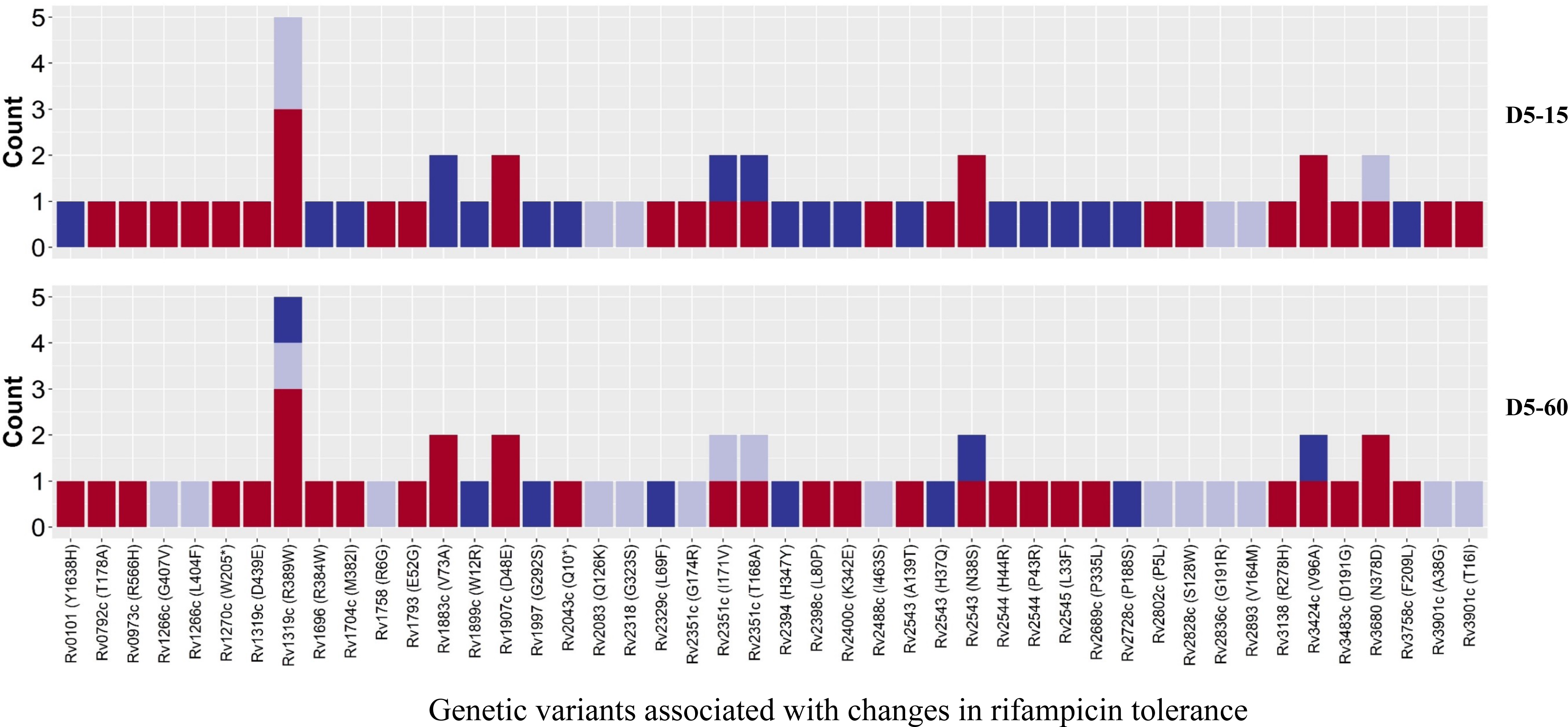
Genetic variants associated with changes in rifampicin tolerance. Non-synonymous single nucleotide polymorphism emerging in pair-wise comparison of longitudinally collected isoniazid resistant *M. tuberculosis* isolates from same patient associated with increase (red), decrease (dark blue) and no change (light violet) in rifampicin tolerance phenotype at 15 and 60 days of incubation (D5-15 and D5-60 respectively). Each count represents a single independent SNP emergence event.

## Discussion

We investigated rifampicin tolerance in a large number of clinical isolates of *M. tuberculosis*. Overall clinical *M. tuberculosis* isolates showed higher levels of rifampicin tolerance than lab isolates as the average survival fraction post-rifampicin treatment decreased only by 90 to 99% over 5 days. We found that levels of rifampicin tolerance are widely distributed among isolates, with some genetically susceptible strains having similar susceptibility to rifampicin-mediated killing as *bona fide* rifampicin-resistant isolates, at least during the 5 days rifampicin exposure of our assay condition. Furthermore, IR isolates were more likely to harbor fast-growing subpopulations with high levels of rifampicin tolerance.

Heterogeneity in regrowth following stress has been linked to a tradeoff between growth fitness and survival^31^, and it is likely that in *M. tuberculosis* such diversification in growth rates among rifampicin-tolerant subpopulations represents such a balance between growth and persistence under antibiotic stress.

We also observed a variation in growth rate in the absence of antibiotic therapy. On average, IR isolates were slower growing than IS isolates, which likely represents a fitness cost due to isoniazid-resistance-causing mutations and strain genetic background^32^. As expected, IS isolates, with slower growth in the absence of drug had a weak association with high levels of rifampicin tolerance at the 15 day time point^30^ (representing rapidly growing recovered cells), whereas both IS and IR isolates with slower growth in the absence of drug were significantly associated with lesser rifampicin survival fraction levels at 60 days– representing slow growing rifampicin tolerant bacilli. These data suggest that slower growth (in absence of drug) in both isoniazid susceptible and resistant isolates, perhaps due to fitness cost of mutations^32^, may be associated with more persister-like tolerant subpopulations.

By contrast, the association between rifampicin MIC and rifampicin tolerance showed a contrasting trend with isoniazid susceptibility. IS isolates showed decreased tolerance with increase in rifampicin MIC, but IR isolates did not show this association. This may indicate higher growth fitness of IR with rifampicin tolerance. Another important finding from our study is the emergence of higher rifampicin tolerance and growth fitness in longitudinal IR isolates under rifampicin treatment selection. This further supports the findings that multiple genetic microvariants may co-exist in patient and rapidly change their proportion under selection from host stresses and antibiotic treatment^33^. We also observed non-synonymous mutations in multiple genes, associated with persistence and host survival enriched with changes in rifampicin tolerance between the longitudinal isolates (supplementary table 3 with references). However, lack of convergent SNPs in the samples may be due to the relatively small sample size, interaction between SNPs and strain background or indication of a larger set of tolerance-related genes that independently affect bacterial growth and antibiotic tolerance^4^.

Our study also reveals novel aspects of rifampicin tolerance associated with isoniazid susceptibility. Rifampicin treatment itself led to the selection of IR *M. tuberculosis* genetic microvariants with high rifampicin tolerance and increased growth fitness in patients. The precise mechanisms underlying these phenotypes will require further investigation, but it is intriguing to note that different *M. tuberculosis* lineages have varying liabilities for development of isoniazid resistance^34^, suggesting that clinical isolates may evolve diverse paths towards phenotypic drug resistance that impact fundamental bacterial physiology and tolerance to other antibiotics.

The wide range of observed rifampicin tolerance, spanning many orders of magnitude confirms findings of experimentally evolved drug tolerance to the laboratory isolate *M. tuberculosis*-H37Rv^10^ and extends our prior findings from a smaller-scale pilot study^27^. Given that almost all rifampicin resistance is via mutations in *rpoB*^35^, our findings suggest that first-line molecular testing for rifampicin susceptibility, which is replacing phenotypic drug susceptibility^36^, may not fully capture response to therapy. It needs to be further validated if these strains that are ‘hyper-tolerant’ to rifampicin are risk factors for poor clinical outcomes in IR-TB^25^.

Given the association of IR with the emergence of rifampicin resistance^26^, our findings suggest a plausible mechanism by which isoniazid resistance, via rifampicin tolerance, acts as a ‘stepping stone’ to rifampicin resistance. The association between IR and rifampicin tolerance only held for fast-growing recovered bacteria. Given the observation that ‘growing’ rifampicin tolerant bacteria are over-represented after initiation of drug therapy in humans due to the specific regulation of *rpoB* in mycobacteria in response to rifampicin exposure^19^, this may represent a divergence between growing and non-replicating persister forms of antibiotic tolerance. A better understanding of which forms of tolerance contribute to clinically relevant response to therapy will be critical for tailoring individualized regimens for TB or improving treatment regimen for IR-TB^37^.

Our study has some limitations. We only assayed rifampicin tolerance under one standard axenic culture condition. It is known that antibiotic tolerance phenotypes vary considerably according to culture conditions^11^, with some phenotypes only emerging *in vitro* with specialized media, e.g. containing odd-chained fatty acids^11^. Secondly, contributors to antibiotic tolerance can be genetic, epigenetic or transient^9–12^, and there is considerable epistasis between genetic variation and antibiotic susceptibility. Our assay will not be able to capture drivers of tolerance that have been lost in the collection, banking, freezing and reviving of the *M. tuberculosis* isolates. Finally, the isolates were from a previous study^25^, and during the study period the old 8-month TB treatment regimen lacked rifampicin in the continuation phase^25^.

This study also reveals interesting aspects like fast and slow growing sub-populations and possible variation in lag-time distribution among clinical *M. tuberculosis* isolates. There can also be different mechanisms of tolerance and persistence among *M. tuberculosis* isolates, detailed investigations are required to further understand these aspects and its clinical relevance.

In conclusion, our study identifies a significant association between isoniazid-resistance and rifampicin tolerance in clinical isolates of *M. tuberculosis*. Our findings have implications for the requirement to consider heterogeneity in bacterial responses to antibiotics and emergence of antibiotic tolerant bacterial genetic microvariants in determining optimal tuberculosis treatment regimens.

## Methods

### Ethical approval

*M. tuberculosis* isolates in this study were a part of collection from a previous study^25^, approved by the Institutional Research Board of Pham Ngoc Thach Hospital as the supervisory institution of the district TB Units (DTUs) in southern Vietnam, Ho Chi Minh City Health Services and the Oxford University Tropical Research Ethics Committee (Oxtrec 030–07).

### Bacterial isolates

242 *M. tuberculosis* isolates, collected for a previous study in Vietnam were used in this study^25^. All the isolates were cultured in the biosafety level-3 laboratory at the Oxford University Clinical Research Unit, Ho Chi Minh city, Vietnam^25^.

### Rifampicin killing assay

Most-probable number-based rifampicin killing assay was done for the clinical *M. tuberculosis* isolates as per the published protocol^27^. *M. tuberculosis* isolates, after single sub-culture from archive, were inoculated in 7H9T medium (Middlebrook 7H9 broth supplemented with 0.2% glycerol, 10% OADC and 0.05% Tween-80) and incubated at 37^0^C until exponential phase with OD_600_ range of 0.4-0.6 is reached. All cultures were homogenized by vortexing for three minutes with sterile glass beads and diluted to the OD_600_ of 0.4. The diluted culture was used for measuring initial viable bacterial number by most probable number (MPN) method, using 96 well plates according to the published protocol^27^.

Briefly the protocol was as follows, a 1 mL aliquot of *M. tuberculosis* culture was harvested, and the cell pellet was washed once. This washed culture was resuspended in 1mL culture and 100 µL was transferred to 96-well plates as an undiluted culture in duplicate for serial dilution. The undiluted culture was used for 10-fold serial dilution up to 10^9^ dilutions in microtiter plates (figure 1B). Immediately, after sampling for initial MPN (day 0), the remaining culture in the tube was treated with rifampicin (Merck-Sigma Aldrich, USA) at a final concentration of 2 µg/mL and incubated. On 2 and 5 days post-rifampicin treatment, viable bacterial number was determined again by MPN method as previously mentioned^27^ (figure 1B). The growth in 96 well plate was recorded as images by the Vizion image system (Thermo Fisher, Scientific Inc, USA) after 15, 30 and 60 days of incubation, beyond 60 days drying of plates were observed (figure 1B). The number of wells with visible bacterial growth was determined by two independent readings from two individuals, discrepancies between the two readings were verified and corrected by a third person reading. MPN value was calculated as mean MPN/mL. The survival fraction at 2 and 5 days post rifampicin treatment was calculated as compared to the initial MPN taken as 100% survival.

### Relative growth difference calculation from MPN number

For calculating relative growth difference of isolates before rifampicin treatment, the log_10_ MPN ratio between 15 and 60 days of incubation were taken to determine the relative proportion of fast and slow growing sub-population. A log_10_ MPN ratio close to 0 indicated less growth heterogeneity in the population, whereas a ratio less than 0 indicated presence of growth heterogeneity due to the presence of fast and slow growing, or heterogeneity in the lag time distribution of sub-populations.

### Drug susceptibility testing

Microtiter drug susceptibility testing was performed using UKMYC6 plates (Thermo Fisher, Scientific Inc·, USA) for determining initial rifampicin and isoniazid phenotypic susceptibility^38^. Briefly, three weeks-old *M. tuberculosis* colonies from Lowenstein-Jensen medium were used to make cellular suspension in 10 mL saline-Tween80 tube with glass beads (Thermo Fisher, Scientific Inc·, USA) and adjusted to 0.5 McFarland units. The suspension is diluted in 7H9 broth (Thermo Fisher, Scientific Inc., USA) and inoculated into 96-well microtiter plate using a semi-automated Sensititre Autoinoculator (Thermo Fisher, Scientific Inc., USA). Plates were sealed with plastic sheet and incubated at 37^0^C for 14 to 21 days. The minimum inhibitory concentration (MIC) was measured by a Sensititre Vizion Digital MIC Viewing system (Thermo Fisher, Scientific Inc., USA) and considered valid if there was growth in the drug free control wells. The clinical resistant cut-off concentrations for isoniazid and rifampicin were 0.1 and 1 µg/mL, respectively.

The IR isolates were also confirmed using BACTEC MGIT 960 SIRE Kit (Becton Dickinson) according to the manufacturer’s instruction in the biosafety level-3 laboratory at the Oxford University Clinical Research Unit^25^. Phenotypic DST was done for streptomycin (1.0 µg/mL), isoniazid (0.1 µg/mL), rifampicin (1.0 µg/mL) and ethambutol (5.0 µg/mL)^25^. Whole genome sequence data was available for the isolates from previously published study^26^ and the Mykrobe predictor TB software platform was used for genotypic antibiotic susceptibility determination of *M. tuberculosis* isolates^39^.

### Statistical analysis

MDK90 values, and its credible interval was estimated using a linear mixed effect model with a Bayesian approach (brm function, brms package).We used the linear mixed effect model for survival analysis as the data consists of repeated measurements at specific time points. For the linear mixed effect model with the bacterial strains as a random effect, we use the Bayesian approach with non-informative priors, which is equivalent to the frequentist approach. The fixed effect relates to the explanatory variable we are utilizing to predict the outcome. Specifically, our outcome measure is the log_10_ survival fraction. The explanatory variables encompass isoniazid susceptibility (categorized as isoniazid susceptible or resistant), the day of sample collection (0, 2, and 5 days), and the duration of incubation (15, and 60 days).

Wilcoxon rank-sum test (stat_compare_means function, ggpubr package) was used to test the null hypothesis that the IS and IR groups have the same continuous distribution, as it is a non-parametric test that does not require a strong assumption about the normality of the distribution of the variable. Chi-Square test (odds ratio function, epitools package) was used to determine if there is a significant relationship between IR and rifampicin tolerance. Cochran Armitage test (CochranArmitageTest function, DescTools package) was performed to test for trend in IR proportion across the levels of rifampicin tolerance. Linear regression (lm function, stats package) was used to evaluate the correlation between rifampicin survival fraction and relative growth.

Statistical analyses and graphs were plotted using R, version 4·0·1^40^ and p-values of ≤0·05 were considered statistically significant.

### MDK_90_, _99_ and _99.99_ calculation

In addition to MDK90 calculated by linear mixed effect model, we also determined the MDK values at 90, 99 and 99.99% reduction in survival fractions for all the *M. tuberculosis* isolates by the following method. The log_10_ MPN values at Day0, Day 2, and Day 5 were used to calculate the respective MDK time for 90%, 99%, and 99.99% reduction in fraction of survival. The calculation of MDK time for individual isolate was based on modelling kill curve as two similar triangles and using the basic proportionality theorem as shown in the flow chart (Supplementary figure 7) to determine the different length of X-axis (Days post rifampicin treatment) corresponding to decline in survival fraction in Y-axis for each MDK time (MDK_90_, _99_ and _99.99_).

For example, in case of MDK90, Y0 (MPN number at day 0), Y2 ((MPN number at day 2), and Y5 ((MPN number at day 5).

First condition tested is, if 90% reduction in survival fraction happened before or at day 2 by checking if log_10_ MPN number at day 2 is less than or equal to 90% reduction as compared to Y0. If the condition is true then the MDK is calculated as x-axis length DF in the two similar triangles modelled in A (triangles ACB and AFD) and corresponding formula for X given below. If the first condition is false then two similar triangles are modelled as in B (triangles ABC and DEC) and X is calculated as 5 – EC. Similarly, for MDK_99_ and MDK_99.99_ time are calculated by applying the condition for 99% and 99.99% reduction in survival fraction.

### Single nucleotide polymorphism difference between longitudinal isoniazid-resistant isolates with differences in rifampicin tolerance

We used whole genome sequence data and genetic variants analysis previously published for identifying non-synonymous single nucleotide polymorphisms (SNPs) emerging in longitudinal isolates from same patients associated with changes in rifampicin tolerance between the isolates^26^.

## Acknowledgements

We acknowledge funding from the Wellcome Trust Intermediate Fellowship in Public Health and Tropical Medicine to NTTT (206724/Z/17/Z), the Wellcome Trust Investigator Award (207487/C/17/Z) and NIAID award (R21AI169005) to BJ and Wellcome Trust Major Overseas Program Funding to GT (106680/B/14/Z). We acknowledge Prof. Rosalind Allen (Professor for Theoretical Microbial Ecology at Friedrich Schiller University of Jena), for reading the manuscript and suggestions.

## References

1. WHO. Global tuberculosis report. https://www.who.int/publications/i/item/9789240037021, 2021.

2. Colangeli R, Jedrey H, Kim S, et al. Bacterial Factors That Predict Relapse after Tuberculosis Therapy. N Engl J Med 2018; 379(9): 823–33.

3. Kwan BW, Valenta JA, Benedik MJ, Wood TK. Arrested protein synthesis increases persister-like cell formation. Antimicrob Agents Chemother 2013; 57(3): 1468–73.

4. Brauner A, Fridman O, Gefen O, Balaban NQ. Distinguishing between resistance, tolerance and persistence to antibiotic treatment. Nat Rev Microbiol 2016; 14(5): 320–30.

5. Ronneau S, Hill PW, Helaine S. Antibiotic persistence and tolerance: not just one and the same. Curr Opin Microbiol 2021; 64: 76–81.

6. Liu Y, Tan S, Huang L, et al. Immune activation of the host cell induces drug tolerance in *Mycobacterium tuberculosis* both in vitro and in vivo. J Exp Med 2016; 213(5): 809–25.

7. Mishra R, Yadav V, Guha M, Singh A. Heterogeneous Host-Pathogen Encounters Coordinate Antibiotic Resilience in *Mycobacterium tuberculosis*. Trends Microbiol 2021; 29(7): 606–20.

8. Gordhan BG, Peters JS, McIvor A, et al. Detection of differentially culturable tubercle bacteria in sputum using mycobacterial culture filtrates. Sci Rep 2021; 11(1): 6493.

9. Su HW, Zhu JH, Li H, et al. The essential mycobacterial amidotransferase GatCAB is a modulator of specific translational fidelity. Nat Microbiol 2016; 1(11): 16147.

10. Torrey HL, Keren I, Via LE, Lee JS, Lewis K. High Persister Mutants in *Mycobacterium tuberculosis*. PLoS One 2016; 11(5): e0155127.

11. Hicks ND, Yang J, Zhang X, et al. Clinically prevalent mutations in *Mycobacterium tuberculosis* alter propionate metabolism and mediate multidrug tolerance. Nat Microbiol 2018; 3(9): 1032–42.

12. Wang BW, Zhu JH, Javid B. Clinically relevant mutations in mycobacterial LepA cause rifampicin-specific phenotypic resistance. Sci Rep 2020; 10(1): 8402.

13. Lee JJ, Lee SK, Song N, et al. Transient drug-tolerance and permanent drug-resistance rely on the trehalose-catalytic shift in *Mycobacterium tuberculosis*. Nat Commun 2019; 10(1): 2928.

14. Imperial MZ, Nahid P, Phillips PPJ, et al. A patient-level pooled analysis of treatment-shortening regimens for drug-susceptible pulmonary tuberculosis. Nat Med 2018; 24(11): 1708–15.

15. Grobbelaar M, Louw GE, Sampson SL, van Helden PD, Donald PR, Warren RM. Evolution of rifampicin treatment for tuberculosis. Infect Genet Evol 2019; 74: 103937.

16. Adams RA, Leon G, Miller NM, et al. Rifamycin antibiotics and the mechanisms of their failure. J Antibiot (Tokyo) 2021; 74(11): 786–98.

17. Adams KN, Takaki K, Connolly LE, et al. Drug tolerance in replicating mycobacteria mediated by a macrophage-induced efflux mechanism. Cell 2011; 145(1): 39–53.

18. Javid B, Sorrentino F, Toosky M, et al. Mycobacterial mistranslation is necessary and sufficient for rifampicin phenotypic resistance. Proc Natl Acad Sci U S A 2014; 111(3): 1132–7.

19. Zhu JH, Wang BW, Pan M, Zeng YN, Rego H, Javid B. Rifampicin can induce antibiotic tolerance in mycobacteria via paradoxical changes in rpoB transcription. Nat Commun 2018; 9(1): 4218.

20. Vijay S, Nair RR, Sharan D, et al. Mycobacterial Cultures Contain Cell Size and Density Specific Sub-populations of Cells with Significant Differential Susceptibility to Antibiotics, Oxidative and Nitrite Stress. Front Microbiol 2017; 8: 463.

21. Aldridge BB, Fernandez-Suarez M, Heller D, et al. Asymmetry and aging of mycobacterial cells lead to variable growth and antibiotic susceptibility. Science 2012; 335(6064): 100–4.

22. Rego EH, Audette RE, Rubin EJ. Deletion of a mycobacterial divisome factor collapses single-cell phenotypic heterogeneity. Nature 2017; 546(7656): 153–7.

23. Mishra R, Kohli S, Malhotra N, et al. Targeting redox heterogeneity to counteract drug tolerance in replicating *Mycobacterium tuberculosis*. Sci Transl Med 2019; 11(518).

24. Saito K, Warrier T, Somersan-Karakaya S, et al. Rifamycin action on RNA polymerase in antibiotic-tolerant *Mycobacterium tuberculosis* results in differentially detectable populations. Proc Natl Acad Sci U S A 2017; 114(24): E4832–E40.

25. Thai PVK, Ha DTM, Hanh NT, et al. Bacterial risk factors for treatment failure and relapse among patients with isoniazid resistant tuberculosis. BMC Infect Dis 2018; 18(1): 112.

26. Srinivasan V, Ha VTN, Vinh DN, et al. Sources of Multidrug Resistance in Patients With Previous Isoniazid-Resistant Tuberculosis Identified Using Whole Genome Sequencing: A Longitudinal Cohort Study. Clin Infect Dis 2020; 71(10): e532–e9.

27. Vijay S, Nhung HN, Bao NLH, et al. Most-Probable-Number-Based Minimum Duration of Killing Assay for Determining the Spectrum of Rifampicin Susceptibility in Clinical *Mycobacterium tuberculosis* Isolates. Antimicrob Agents Chemother 2021; 65(3).

28. Prakash J, Velpandian T, Pande JN, Gupta SK. Serum Rifampicin Levels in Patients with Tuberculosis : Effect of P-Glycoprotein and CYP3A4 Blockers on its Absorption. Clin Drug Investig 2003; 23(7): 463–72.

29. Von Groll A, Martin A, Portaels F, Silva PEAd, Palomino JCJBJoM. Growth kinetics of *Mycobacterium tuberculosis* measured by quantitative resazurin reduction assay: a tool for fitness studies. 2010; 41: 300–3.

30. Pontes MH, Groisman EA. Slow growth determines nonheritable antibiotic resistance in *Salmonella enterica*. Sci Signal 2019; 12(592).

31. Moreno-Gamez S, Kiviet DJ, Vulin C, et al. Wide lag time distributions break a trade-off between reproduction and survival in bacteria. Proc Natl Acad Sci U S A 2020; 117(31): 18729–36.

32. Gagneux S. Fitness cost of drug resistance in *Mycobacterium tuberculosis*. Clin Microbiol Infect 2009; 15 **Suppl 1**: 66–8.

33. Trauner A, Liu Q, Via LE, et al. The within-host population dynamics of Mycobacterium tuberculosis vary with treatment efficacy. Genome Biol 2017; 18(1): 71.

34. Carey AF, Rock JM, Krieger IV, et al. TnSeq of Mycobacterium tuberculosis clinical isolates reveals strain-specific antibiotic liabilities. PLoS Pathog 2018; 14(3): e1006939.

35. Zaw MT, Emran NA, Lin Z. Mutations inside rifampicin-resistance determining region of rpoB gene associated with rifampicin-resistance in *Mycobacterium tuberculosis*. J Infect Public Health 2018; 11(5): 605–10.

36. Macedo R, Nunes A, Portugal I, Duarte S, Vieira L, Gomes JP. Dissecting whole-genome sequencing-based online tools for predicting resistance in *Mycobacterium tuberculosis*: can we use them for clinical decision guidance? Tuberculosis (Edinb) 2018; 110: 44–51.

37. WHO. WHO treatment guidelines for isoniazid-resistant tuberculosis., 2018.

38. Rancoita PMV, Cugnata F, Gibertoni Cruz AL, et al. Validating a 14-Drug Microtiter Plate Containing Bedaquiline and Delamanid for Large-Scale Research Susceptibility Testing of *Mycobacterium tuberculosis*. Antimicrob Agents Chemother 2018; 62(9).

39. Bradley P, Gordon NC, Walker TM, et al. Rapid antibiotic-resistance predictions from genome sequence data for *Staphylococcus aureus* and *Mycobacterium tuberculosis*. Nat Commun 2015; 6: 10063.

40. Team RC. R: A language and environment for statistical computing. R Foundation for Statistical Computing, Vienna, Austria. 2012. 2018.

