## Supplemental material for "Rifampicin tolerance and growth fitness among isoniazid-resistant clinical *Mycobacterium tuberculosis* isolates: an in-vitro longitudinal study"

**Supplementary information**

**Supplementary Figures**

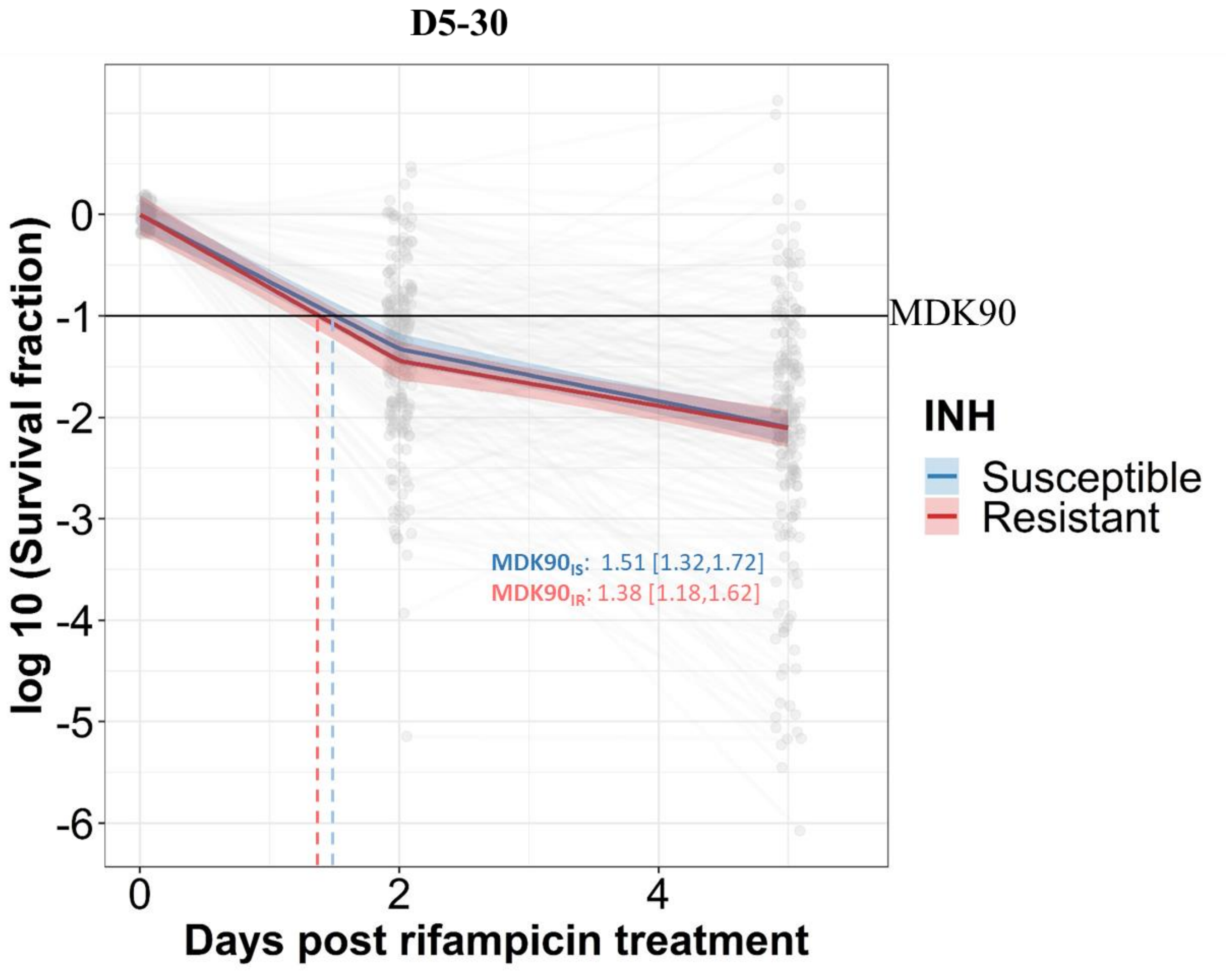

**Supplementary figure 1. Rifampicin survival curve in isoniazid susceptible and resistant clinical *M. tuberculosis* isolates. The bacterial kill curve as measured by $\log_{10}$ survival fraction from data collected at 0, 2 and 5 days of rifampicin treatment followed by incubation for 30 days. Data from individual isolates are shown as the grey dots connected by lines. Estimated mean with 95% credible interval (bold coloured line and colour shaded area respectively) of isoniazid susceptible (IS – blue) and resistant (IR – red) clinical *M. tuberculosis* isolates based on linear mixed effect model implemented in a Bayesian framework. One $\log_{10}$ fold or 90% reduction in survival fraction is indicated (MDK90, black horizontal line) and the mean time duration required for 90% reduction in survival (MDK90, minimum duration of killing time) is indicated by vertical dashed lines with respective colours for IS and IR isolates.**

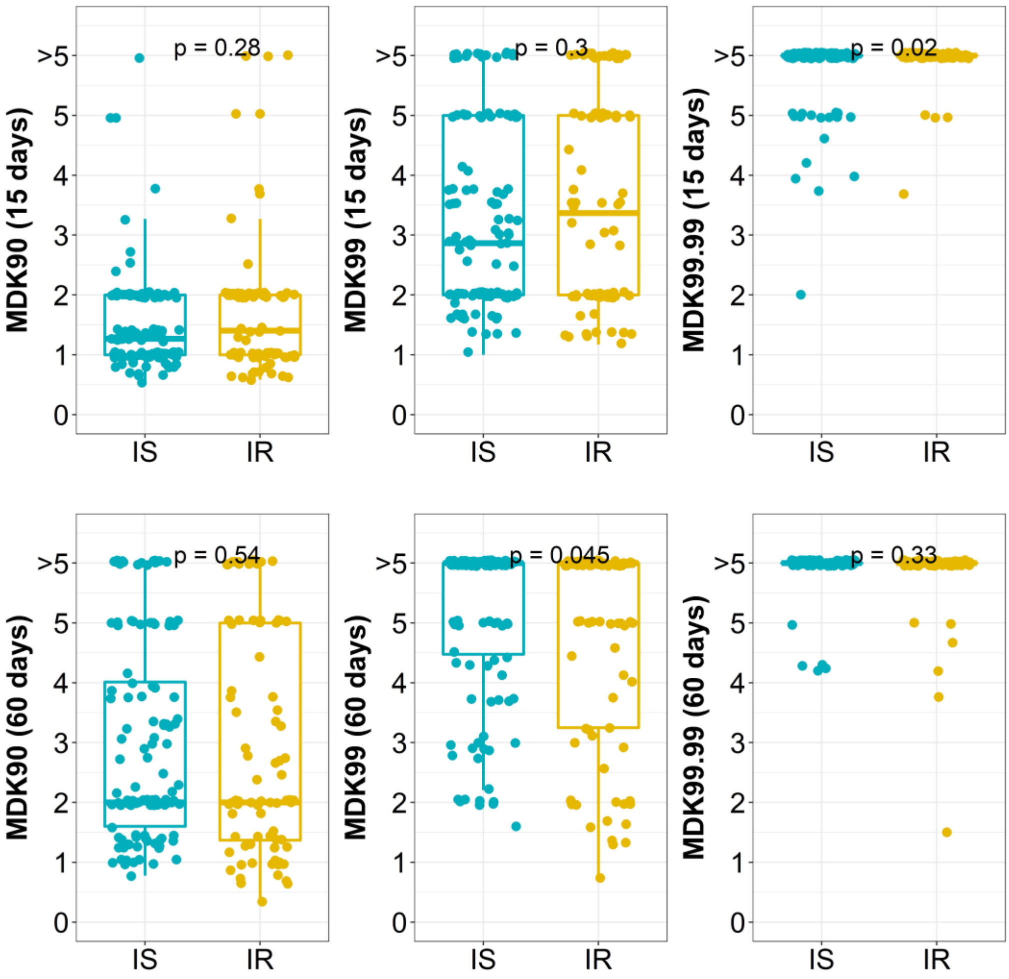

**Supplementary figure 2. Distribution of MDK$_{90}$, $_{99}$ and $_{99.99}$ time (in days) for isoniazid susceptible (IS) and resistant (IR) isolates at 15 and 60 days incubation. Statistical comparisons between IS and IR were made by using Wilcoxon rank-sum test. >5 indicate MDK time above the assay limit.**

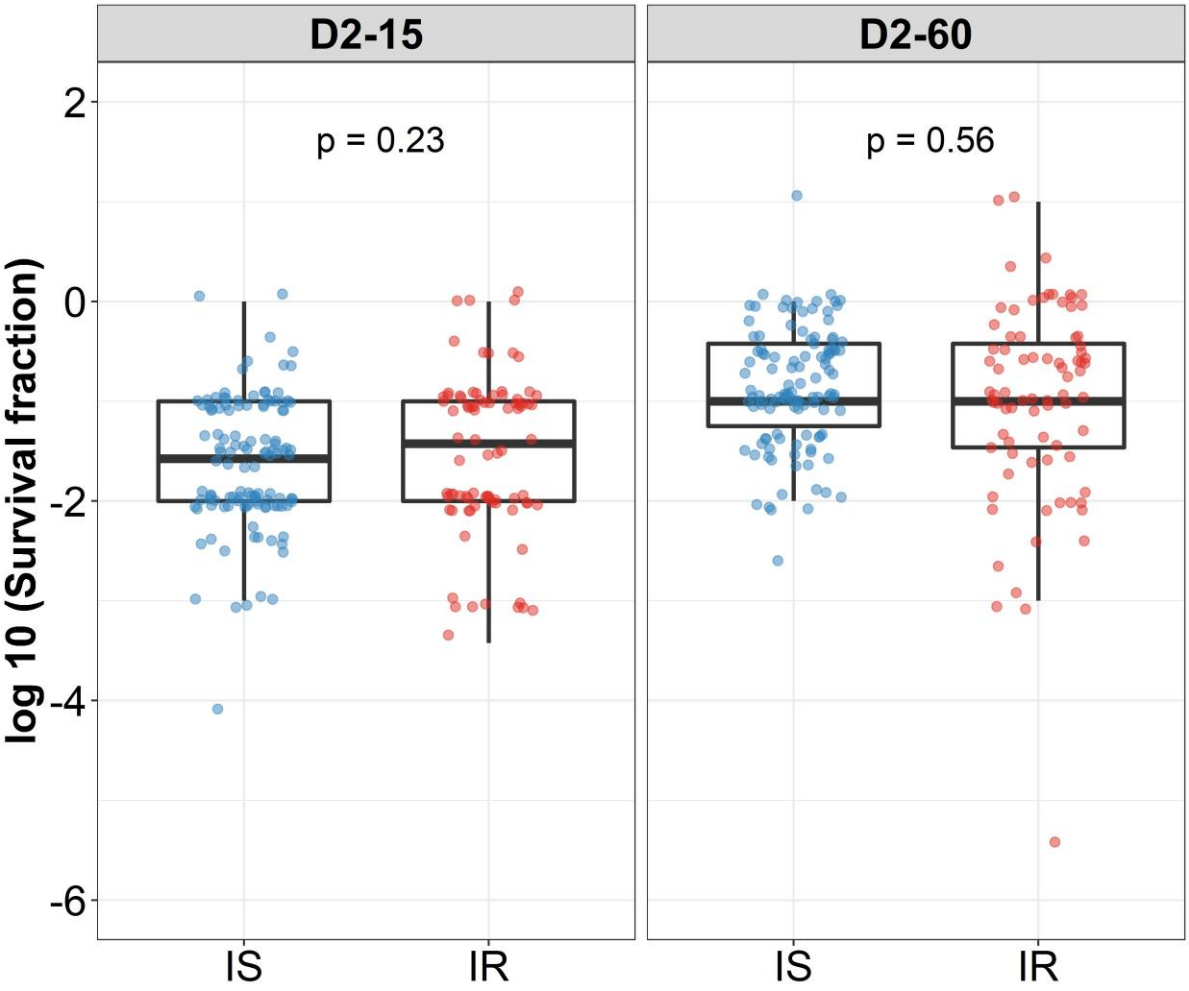

**Supplementary figure 3. $Log_{10}$ survival fraction distribution in isoniazid susceptible (IS) and resistant (IR) clinical *M. tuberculosis* isolates post 2 days of rifampicin treatment at 15 (D2-15) and 60 (D2-60) days of incubation. Statistical comparisons between IS and IR were made by using Wilcoxon rank-sum test.**

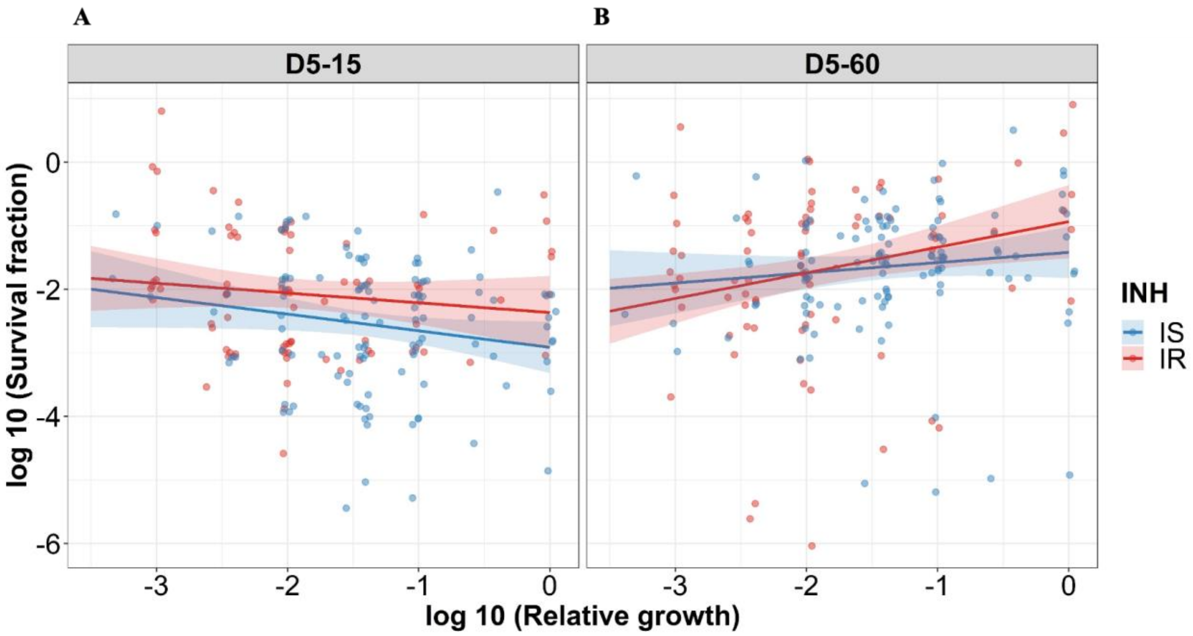

**Supplementary figure 4. Correlating rifampicin survival fraction with before treatment relative growth of clinical *M. tuberculosis* isolates with outliers included. $Log_{10}$ survival fraction at 5 days of rifampicin treatment as determined at 15 days incubation (A) and for isoniazid susceptible (IS, blue dots) and resistant (IR, red dots) isolates respectively, and at 60 days of incubation (B), and for IS (blue dots) and IR (red dots) isolates respectively, correlated with the $log_{10}$ relative growth determined before rifampicin treatment for clinical *M. tuberculosis* isolates. Coefficients of linear regression for (A) IS = -0.26 [-0.52, -0.0017], *P* = 0.051; IR = -0.15 [-0.43, 0.13], *P* = 0.28, and (B) IS = 0.16 [-0.10, 0.42], *P* = 0.23; IR = 0.40 [0.12, 0.68], *P* = 0.005.**

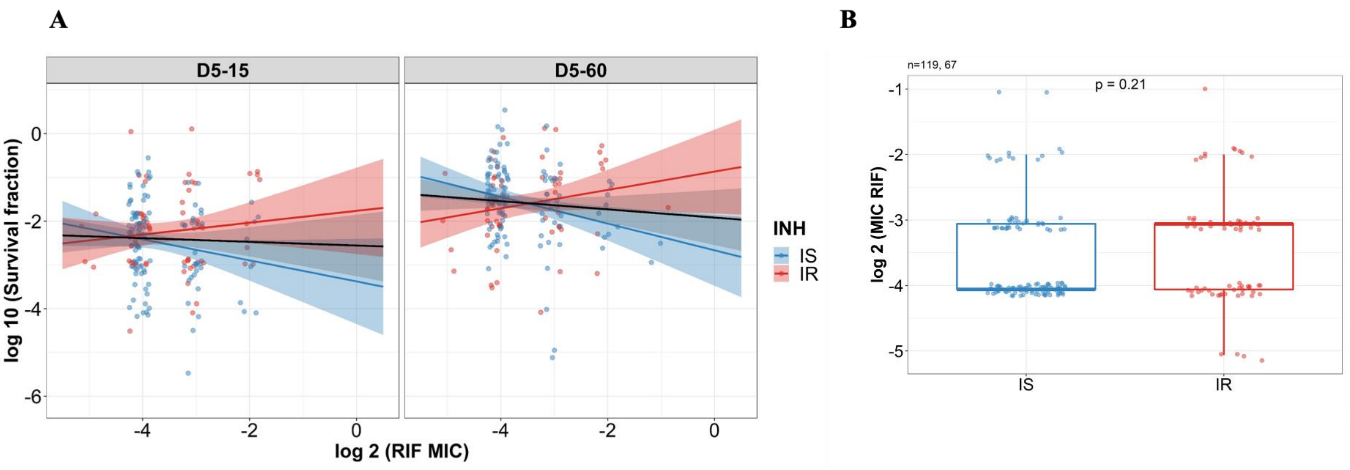

**Supplementary Figure 5. Correlating rifampicin survival fraction with rifampicin MIC of clinical *M. tuberculosis* isolates. (A) $Log_{10}$ survival fraction at 5 days of rifampicin treatment as determined at 15 days (D5-15) and 60 days of incubation (D5-60) for isoniazid susceptible (IS, blue dots) and resistant (IR, red dots) isolates respectively, correlated with the rifampicin MIC of clinical *M. tuberculosis* isolates (MIC range 0.03, 0.06, 0.12, 0.25, 0.5 µg/mL converted to $log_2$). Coefficients of linear regression for (D5-15) IS = -0.24 [-0.50, 0.022], *P* = 0.073; IR = 0.14 [-0.14, 0.41], *P* = 0.33, and (D5-60) IS = -0.31 [-0.53, -0.083], *P* = 0.007; IR = 0.21 [-0.057, 0.48], *P* = 0.12. Black line indicate overall trend of MIC distribution for all isolates. (B) Rifampicin MIC distribution between IS (n=119) and IR (n=67) clinical *M. tuberculosis* isolates. Statistical comparisons between IS and IR were made by using Wilcoxon rank-sum test.**

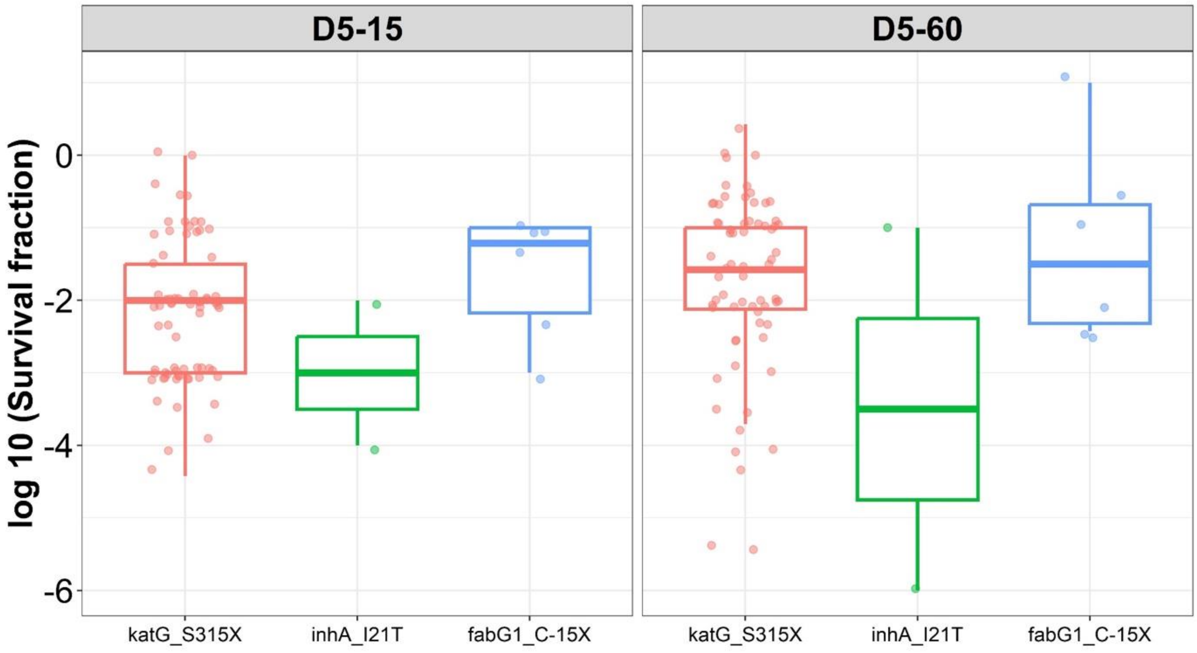

**Supplementary Figure 6. Rifampicin tolerance distribution grouped based on isoniazid resistant mutations (katG_S315X, inhA_I21T, and fabG1_C-15X) in *M. tuberculosis* isolates.**

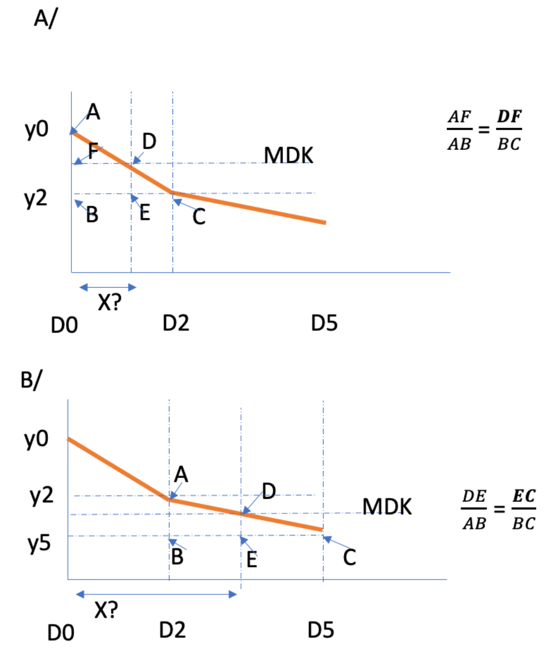

MDK90 calculation
(y in log scale)

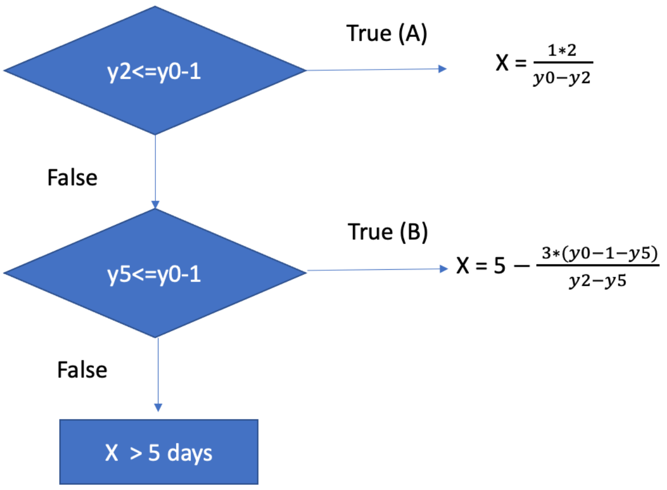

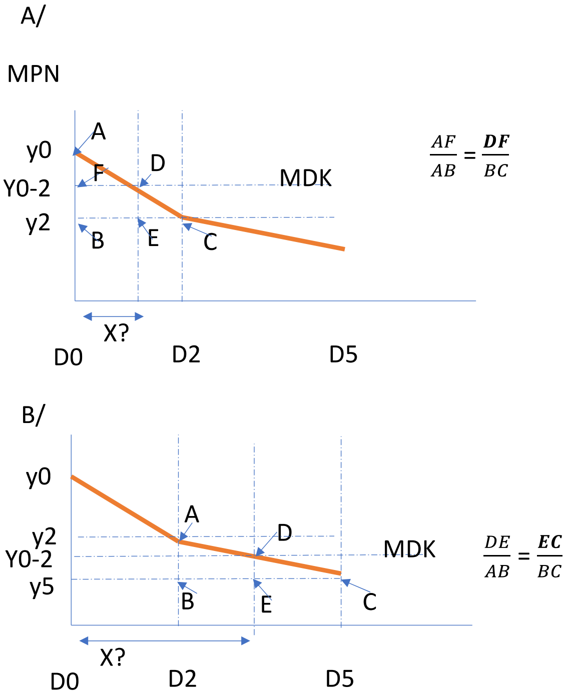

MDK99 calculation

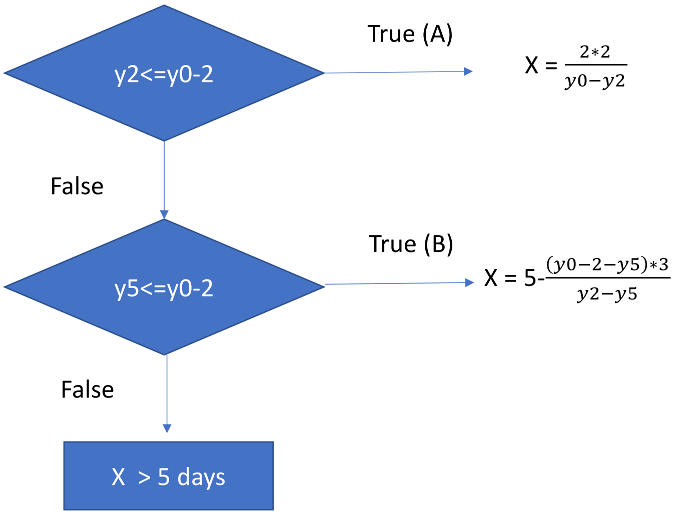

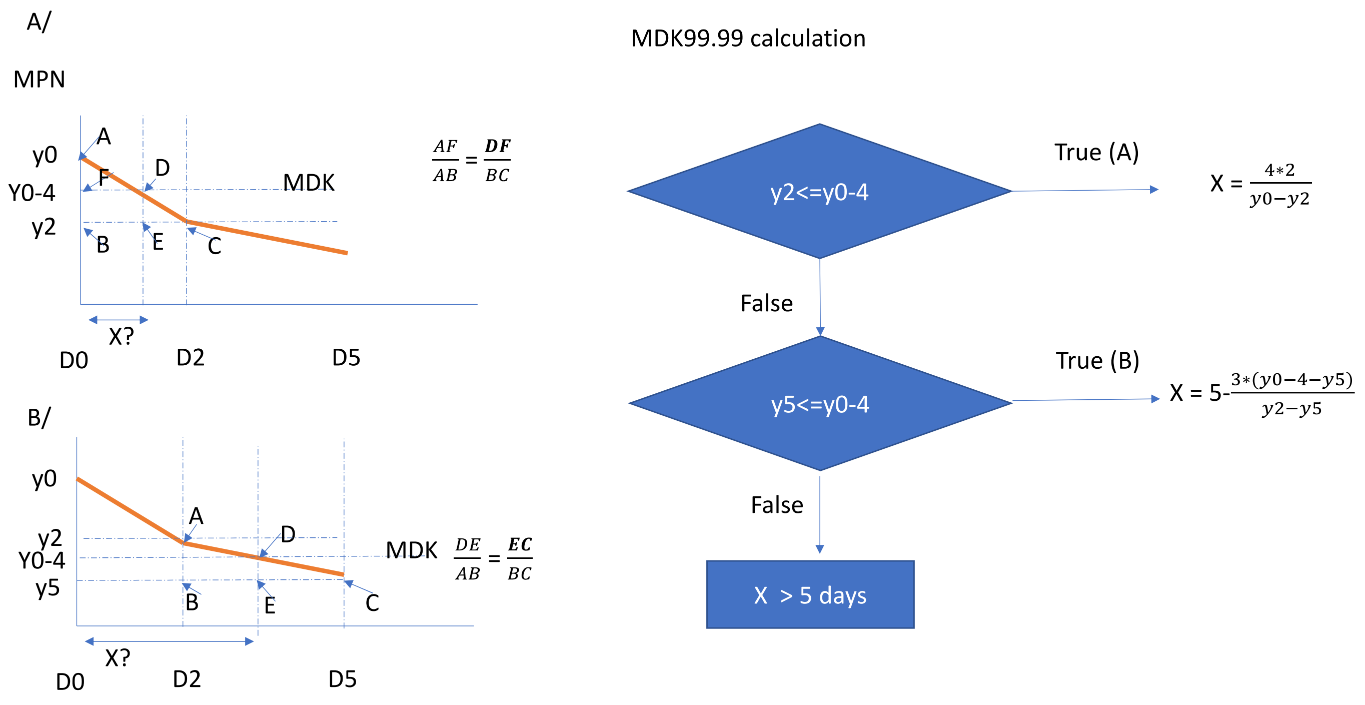

**Supplementary Figure 7. Flow chart for calculating MDK 90, 99 and 99.99 time for clinical *M. tuberculosis* isolates.**

### Supplementary Tables

Supplementary table 1. Rifampicin 2 µg/mL as times of MIC for *M. tuberculosis* isolates.

| Number of *M. tuberculosis* isolates | Rifampicin MIC | Rifampicin 2 µg/mL as times of MIC |
|---|---|---|
| 4 | 0.03 | 66.6 |
| 104 | 0.06 | 33.3 |
| 55 | 0.12 | 16.6 |
| 20 | 0.25 | 8.0 |
| 3 | 0.5 | 4.0 |

Supplementary table 2. Emerging non-synonymous SNPs between longitudinal isoniazid resistant isolates from same patient.

| Patient number | trend_group_D 5_15 | trend_group_D 5_60 | *M. tb* isolate collection (in months) | Compari-sion between two isolates | SNP differe -nce | isolate 1 | isolate 2 | amino_ acid change | Locus_gene | product | gene | locus_tag | Repetiti ve-e_region |
|---|---|---|---|---|---|---|---|---|---|---|---|---|---|
| 1 | Increase | Increase | 0M, 1M, 8M | 0M, 1M | 0 | NA | ad_ratio :0.9 | E52G | Rv1793 (esxN) | ESAT-6 like protein EsxN | esxN | Rv1793 | FALSE |
| 1 | Increase | Increase | 0M, 1M, 8M | 0M, 1M | 0 | NA | ad_ratio :0.9 | N38S | Rv2543 (lppA) | lipoprotein LppA | lppA | Rv2543 | FALSE |
| 1 | Increase | Increase | 0M, 1M, 8M | 0M, 1M | 0 | NA | ad_ratio :0.9 | R278H | Rv3138 (pflA) | pyruvate formate lyase activating protein PflA | pflA | Rv3138 | FALSE |
| 1 | Increase | Increase | 0M, 1M, 8M | 0M, 1M | 0 | ad_ratio: 0.9 | NA | R389W | Rv1319c (NA) | adenylate cyclase | NA | Rv1319c | FALSE |
| 2 | Increase | Decrease | 0M, 1M, 12M | 0M, 1M | 0 | NA | ad_ratio :0.9 | H37Q | Rv2543 (lppA) | lipoprotein LppA | lppA | Rv2543 | FALSE |
| 2 | Increase | Decrease | 0M, 1M, 12M | 0M, 1M | 0 | NA | ad_ratio :0.9 | N38S | Rv2543 (lppA) | lipoprotein LppA | lppA | Rv2543 | FALSE |
| 3 | Increase | Unchange | 0M, 1M | 0M, 1M | 3 | NA | ad_ratio :0.9 | G407V | Rv1266c (pknH) | serine/threo nine-protein kinase PknH | pknH | Rv1266c | FALSE |
| 3 | Increase | Unchange | 0M, 1M | 0M, 1M | 3 | NA | ad_ratio :0.9 | L404F | Rv1266c (pknH) | serine/threo nine-protein kinase PknH | pknH | Rv1266c | FALSE |
| 3 | Increase | Unchange | 0M, 1M | 0M, 1M | 3 | NA | ad_ratio :0.9 | G174R | Rv2351c (plcA) | membrane-associated phospholipa se A | plcA | Rv2351c | FALSE |
| 3 | Increase | Unchange | 0M, 1M | 0M, 1M | 3 | NA | ad_ratio :0.9 | I171V | Rv2351c (plcA) | membrane-associated phospholipa se A | plcA | Rv2351c | FALSE |
| 3 | Increase | Unchange | 0M, 1M | 0M, 1M | 3 | NA | ad_ratio :0.9 | T168A | Rv2351c (plcA) | membrane-associated phospholipa se A | plcA | Rv2351c | FALSE |
| 3 | Increase | Unchange | 0M, 1M | 0M, 1M | 3 | NA | PASS | P5L | Rv2802c (NA) | arginine/hy pothetical protein | NA | Rv2802c | FALSE |
| 3 | Increase | Unchange | 0M, 1M | 0M, 1M | 3 | NA | ad_ratio :0.9 | A38G | Rv3901c (NA) | membrane protein | NA | Rv3901c | FALSE |
| 3 | Increase | Unchange | 0M, 1M | 0M, 1M | 3 | NA | ad_ratio :0.9 | T16I | Rv3901c (NA) | membrane protein | NA | Rv3901c | FALSE |
| 3 | Increase | Unchange | 0M, 1M | 0M, 1M | 3 | ad_ratio: 0.9 | NA | R6G | Rv1758 (cut1) | cutinase | cut1 | Rv1758 | FALSE |
| 3 | Increase | Unchange | 0M, 1M | 0M, 1M | 3 | PASS | NA | I463S | Rv2488c (NA) | LuxR family transcriptio nal regulator | NA | Rv2488c | FALSE |
| 3 | Increase | Unchange | 0M, 1M | 0M, 1M | 3 | PASS | NA | S128W | Rv2828c (NA) | hypothetical protein | NA | Rv2828c | FALSE |
| 4 | Increase | Increase | 0M, 1M, 2M, 12M, | 0M, 12M | 0 | NA | ad_ratio :0.9 | T178A | Rv0792c (NA) | transcriptio nal regulator | NA | Rv0792c | FALSE |
| 4 | Increase | Increase | 0M, 1M, 2M, 12M, | 0M, 12M | 0 | NA | ad_ratio :0.9 | D48E | Rv1907c (NA) | hypothetical protein | NA | Rv1907c | FALSE |
| 4 | Increase | Increase | 0M, 1M, 2M, 12M, | 0M, 12M | 0 | NA | ad_ratio :0.9 | V96A | Rv3424c (NA) | hypothetical protein | NA | Rv3424c | FALSE |
| 4 | Increase | Increase | 0M, 1M, 2M, 12M, | 0M, 12M | 0 | ad_ratio: 0.9 | NA | R389W | Rv1319c (NA) | adenylate cyclase | NA | Rv1319c | FALSE |
| 4 | Increase | Increase | 0M, 1M, 2M, 12M, | 0M, 12M | 0 | ad_ratio: 0.9 | NA | N378D | Rv3680 (NA) | anion transporter ATPase | NA | Rv3680 | FALSE |
| 5 | Increase | Decrease | 0M, 24M | 0M, 24M | 3 | NA | PASS | L69F | Rv2329c (narK1) | nitrate/nitrit e transporter | narK 1 | Rv2329c | FALSE |
| 5 | Increase | Decrease | 0M, 24M | 0M, 24M | 3 | ad_ratio: 0.9 | NA | R389W | Rv1319c (NA) | adenylate cyclase | NA | Rv1319c | FALSE |
| 7 | Increase | Decrease | 0M, 1M, 18M | 0M, 1M | 0 | NA | ad_ratio :0.9 | V96A | Rv3424c (NA) | hypothetical protein | NA | Rv3424c | FALSE |
| 8 | Increase | Increase | 0M, 2M | 0M, 2M | 0 | NA | ad_ratio :0.9 | D48E | Rv1907c (NA) | hypothetical protein | NA | Rv1907c | FALSE |
| 9 | Increase | Increase | 0M, 1M, 12M | 0M, 12M | 3 | NA | PASS | R566H | Rv0973c (accA2) | acetyl/propi onyl-CoA carboxylase subuit alpha | accA 2 | Rv0973c | FALSE |

| | | | | | | | | | | | | | |
|---|---|---|---|---|---|---|---|---|---|---|---|---|---|
| 9 | Increase | Increase | 0M, 1M, 12M | 0M, 12M | 3 | NA | PASS | W205* | Rv1270c (lprA) | lipoprotein LprA | lprA | Rv1270c | FALSE |
| 9 | Increase | Increase | 0M, 1M, 12M | 0M, 12M | 3 | NA | ad_ratio :0.9 | D439E | Rv1319c (NA) | adenylate cyclase | NA | Rv1319c | FALSE |
| 9 | Increase | Increase | 0M, 1M, 12M | 0M, 12M | 3 | NA | PASS | D191G | Rv3483c (NA) | hypothetical protein | NA | Rv3483c | FALSE |
| 12 | Unchange | Increase | 0M, 8M | 0M, 8M | 1 | ad_ratio: 0.9 | NA | R389W | Rv1319c (NA) | adenylate cyclase | NA | Rv1319c | FALSE |
| 12 | Unchange | Increase | 0M, 8M | 0M, 8M | 1 | ad_ratio: 0.9 | NA | N378D | Rv3680 (NA) | anion transporter ATPase | NA | Rv3680 | FALSE |
| 13 | Unchange | Unchange | 0M, 12M | 0M, 12M | 3 | NA | PASS | Q126K | Rv2083 (NA) | hypothetical protein | NA | Rv2083 | FALSE |
| 13 | Unchange | Unchange | 0M, 12M | 0M, 12M | 3 | ad_ratio: 0.9 | NA | R389W | Rv1319c (NA) | adenylate cyclase | NA | Rv1319c | FALSE |
| 13 | Unchange | Unchange | 0M, 12M | 0M, 12M | 3 | ad_ratio: 0.9 | NA | G323S | Rv2318 (uspC) | sugar ABC transporter substrate-binding lipoprotein UspC | uspC | Rv2318 | FALSE |
| 13 | Unchange | Unchange | 0M, 12M | 0M, 12M | 3 | PASS | NA | G191R | Rv2836c (dinF) | DNA-damage-inducible protein DinF | dinF | Rv2836c | FALSE |
| 13 | Unchange | Unchange | 0M, 12M | 0M, 12M | 3 | PASS | NA | V164M | Rv2893 (NA) | oxidoreductase | NA | Rv2893 | FALSE |
| 14 | Decrease | Decrease | 0M, 18M | 0M, 18M | 3 | NA | ad_ratio :0.9 | G292S | Rv1997 (ctpF) | cation transporter ATPase F | ctpF | Rv1997 | FALSE |
| 14 | Decrease | Decrease | 0M, 18M | 0M, 18M | 3 | NA | PASS | H347Y | Rv2394 (ggtB) | gamma-glutamyltranspeptidase precursor GgtB | ggtB | Rv2394 | FALSE |
| 14 | Decrease | Decrease | 0M, 18M | 0M, 18M | 3 | NA | PASS | P188S | Rv2728c (NA) | hypothetical protein | NA | Rv2728c | FALSE |
| 14 | Decrease | Decrease | 0M, 18M | 0M, 18M | 3 | PASS | NA | W12R | Rv1899c (lppD) | lipoprotein LppD | lppD | Rv1899c | FALSE |
| 16 | Decrease | Increase | 0M, 5M | 0M, 5M | 2 | NA | PASS | M382I | Rv1704c (cycA) | D-serine/alanine/glycine transporter protein CycA | cycA | Rv1704c | FALSE |
| 16 | Decrease | Increase | 0M, 5M | 0M, 5M | 2 | NA | ad_ratio :0.9 | V73A | Rv1883c (NA) | hypothetical protein | NA | Rv1883c | FALSE |
| 16 | Decrease | Increase | 0M, 5M | 0M, 5M | 2 | ad_ratio: 0.9 | NA | I171V | Rv2351c (plcA) | membrane-associated phospholipase A | plcA | Rv2351c | FALSE |
| 16 | Decrease | Increase | 0M, 5M | 0M, 5M | 2 | ad_ratio: 0.9 | NA | T168A | Rv2351c (plcA) | membrane-associated phospholipase A | plcA | Rv2351c | FALSE |
| 16 | Decrease | Increase | 0M, 5M | 0M, 5M | 2 | ad_ratio: 0.9 | NA | A139T | Rv2543 (lppA) | lipoprotein LppA | lppA | Rv2543 | FALSE |
| 17 | Decrease | Increase | 1M, 8M | 1M, 8M | 11 | NA | PASS | Y1638H | Rv0101 (nrp) | peptide synthetase Nrp | nrp | Rv0101 | FALSE |
| 17 | Decrease | Increase | 1M, 8M | 1M, 8M | 11 | NA | PASS | R384W | Rv1696 (recN) | DNA repair protein RecN | recN | Rv1696 | FALSE |
| 17 | Decrease | Increase | 1M, 8M | 1M, 8M | 11 | NA | PASS | Q10* | Rv2043c (pncA) | pyrazinamidase/nicotinamidase PncA | pncA | Rv2043c | FALSE |
| 17 | Decrease | Increase | 1M, 8M | 1M, 8M | 11 | NA | PASS | K342E | Rv2400c (subI) | sulfate ABC transporter substrate-binding lipoprotein SubI | subI | Rv2400c | FALSE |
| 17 | Decrease | Increase | 1M, 8M | 1M, 8M | 11 | NA | PASS | P43R | Rv2544 (lppB) | lipoprotein LppB | lppB | Rv2544 | FALSE |
| 17 | Decrease | Increase | 1M, 8M | 1M, 8M | 11 | NA | PASS | H44R | Rv2544 (lppB) | lipoprotein LppB | lppB | Rv2544 | FALSE |
| 17 | Decrease | Increase | 1M, 8M | 1M, 8M | 11 | NA | ad_ratio :0.9 | L33F | Rv2545 (vapB18) | antitoxin VapB18 | vapB 18 | Rv2545 | FALSE |
| 17 | Decrease | Increase | 1M, 8M | 1M, 8M | 11 | NA | PASS | P335L | Rv2689c (NA) | hypothetical protein | NA | Rv2689c | FALSE |
| 17 | Decrease | Increase | 1M, 8M | 1M, 8M | 11 | ad_ratio: 0.9 | NA | V73A | Rv1883c (NA) | hypothetical protein | NA | Rv1883c | FALSE |
| 17 | Decrease | Increase | 1M, 8M | 1M, 8M | 11 | PASS | NA | L80P | Rv2398c (cysW) | sulfate ABC transporter permease CysW | cysW | Rv2398c | FALSE |
| 18 | Decrease | Increase | 0M, 12M, 18M | 0M, 18M | 3 | NA | PASS | F209L | Rv3758c (proV) | glycine betaine/carnitine/choline/L-proline ABC | proV | Rv3758c | FALSE |

| | | | | | | | | | | |
|---|---|---|---|---|---|---|---|---|---|---|
| | | | | | | | | | | transporter ATP-binding protein ProV |

NA - Wild-type

ad_ratio:0.9 - Emerging Non-synonymous SNP in reads below 90% threshold.

Pass - Emerging Non-synonymous SNP in reads above 90% threshold.

Note: SNPs difference include both synonymous and nonsynonymous variants (only Pass).

Supplementary table 3. Genes with emergence of non-synonymous mutations associated with changes in rifampicin tolerance between longitudinal isoniazid resistant *M. tuberculosis* isolates.

| Gene name | Mutation | Gene function | Associated change rifampicin tolerance – D5-15 | Associated change rifampicin tolerance – D5-60 | Reported function related to survival or antibiotic response | Reference |
|---|---|---|---|---|---|---|
| Rv0101 | Y1638H | Probable peptide synthetase nrp | Decrease | Increase | • Down regulated upon rifampicin exposure of MDR-H37Rv<br>• Prevented INH-mediated killing *M. tuberculosis* in Mice | Knegt et. al., 2013<br><br>Dhar and McKinney 2010 |
| Rv0792c | T178A | Probable transcription regulatory protein (gntr family) | Increase | Increase | Gene upregulated in *M. tuberculosis* persisters. | Keren et. al., 2011 |
| Rv0973c | R566H | Lipid metabolism | Increase | Increase | Gene downregulated upon HigB toxin expression in *M. tuberculosis* | Schuessler et. al., 2013 |
| Rv1266c | G407V | *pknH*, Serine/threonine protein kinase | Increase | No change | Interaction partner of FtsB in *M. tuberculosis,* regulation of cell division during persistence. | Wang et. al., 2019 |
| | L404F | | Increase | No change | | |
| Rv1270c | W205* | *lprA*, Cell wall and cell processes | Increase | Increase | Gene upregulated upon *sigF* induction | Williams et. al., 2007 |
| Rv1319c | D439E | Putative adenylate cyclase, regulation of cellular metabolism | Increase | Increase | Gene displaying high within-host genetic diversity. | Nimmo et. al., 2020 |
| | R389W | | Increase (n = 3), No change (n = 2), Decrease (n = 1). | Increase (n = 3), no change (n = 1), decrease (n = 1). | | |
| Rv1696 | R384W | *RccN*, works in DNA repair | Decrease | Increase | DNA repair mechanism in *M. tuberculosis* | Mittal et. al., 2020 |
| Rv1704c | M382I | Alanine transporter CycA | Decrease | Increase | Gene involved in D-cycloserine resistance | Vilcheze 2020 |
| Rv1758 | R6G | | Increase | Decrease | Gene frameshift deletion associated with hypervirulence and enhanced growth in macrophage. | Lam et. al., 2011 |

| | | | | | | |
|---|---|---|---|---|---|---|
| Rv1883c | V73A | | Decrease (n = 2) | Increase (n = 2) | Known mutation associated with IR | Lagutkin et. al., 2022 |
| Rv1899c | W12R | | Decrease | Decrease | No report | |
| Rv1907c | D48E | | Increase (n = 2) | Increase (n = 2) | Gene associated with IR | Lagutkin et. al., 2022 |
| Rv1997 | G292S | *ctpF M. tuberculosis* cation transporter | Decrease | Decrease | Gene regulated by DosR, involved in hypoxia, dormancy regulon and ion transport | Pulido et. al., 2014 |
| Rv2043c | Q10* | *pncA*, pyrazinamidase enzyme | Decrease | Increase | Gene involved in Pyrazinamide resistance | Baddam et. al., 2018 |
| Rv2083 | Q126K | | No change | No change | | |
| Rv2318 | G323S | | No change | No change | | |
| Rv2329c | L69F | *narK1* | Increase | Decrease | Gene overexpressed under nitrogen limitation in *M. tuberculosis* | Williams et. al., 2015 |
| Rv2351c | G174R | *plcA* Membrane-associated phospholipase C | Increase | Decrease | Gene displaying high within-host genetic diversity. | Shockey et. al., 2019 |
| | I171V | | Increase (n = 1) Decrease (n = 1) | Increase (n = 1) No change (n = 1) | | |
| | T168A | | Increase (n = 1) Decrease (n = 1) | Increase (n = 1) No change (n = 1) | | |
| Rv2394 | H347Y | *ggtB*, Oxidative stress proteins | Decrease | Decrease | Gene upregulated in *M. tuberculosis* under oxidative stress induced by sulfamethoxazole | Sarkar et. al., 2018 |
| Rv2398c | L80P | *cysW*, Sulphate transport system permease protein, resuscitation-promoting factor. | Decrease | Increase | Gene upregulated under starvation model of *M. tuberculosis* persistence | Betts et. al., 2002 Gorla et. al., 2018 |
| Rv2400c | K342E | *subI* Sulphate binding precursor | Decrease | Increase | • Gene upregulated under starvation model of *M. tuberculosis* persistence<br>• Possible role in antibiotic tolerance. | Betts et. al., 2002<br>Xu et. al., 2017 |
| Rv2488c | I463S | LuxR family regulator | Increase | Decrease | Gene family involved in *M. tuberculosis* dormancy and virulence | Fang et. al., 2013 |
| Rv2543 | A139T | *IppA* | Decrease | Increase | Regulated by stationary phase sigma factor *sigD* | Calamita et. al., 2005 |
| | H37Q | | Increase | Decrease | | |
| | N38S | | Increase (n = 2) | Increase (n = 1) Decrease | | |

| | | | | | | |
|---|---|---|---|---|---|---|
| | | | | (n = 1) | | |
| Rv2544 | H44R | Membrane lipoprotein | Decrease | Decrease | | Fleischmann et. al., 2002 |
| | P43R | | Increase | Increase | | |
| Rv2545 | L33F | *vapBC* Toxin-antitoxin modules | Decrease | Increase | Regulate bacterial growth arrest. | Ahidjo et. al., 2011 |
| Rv2689c | P335L | | Decrease | Increase | Gene with mutation during *M. tuberculosis* latent infection | Colangeli et. al., 2014 |
| Rv2728c | P188S | Conserved alanine rich protein | Decrease | Decrease | | Li et. al., 2015 |
| Rv2836c | G191R | *dinF* | No change | No change | Gene associated with DNA repair and upregulated in pulmonary TB | Rachman et. al., 2006 |
| Rv2893 | V164M | Similar to alkanal monooxygenase alpha chain | No change | No change | Gene repressed in acid stress | Fisher et. al., 2002 |
| Rv3138 | R278H | Pyruvate–formate–lyase activating protein | Increase | Increase | Gene upregulated in *M. tuberculosis* stationary phase | Hampshire et. al., 2004 |
| Rv3424c | V96A | | Increase (n = 2) | Increase (n = 1) Decrease (n = 1) | Gene displaying high within-host genetic diversity. | Nimmo et. al., 2020 |
| Rv3483c | D191G | | Increase | Increase | MmpL3 (mycolic acid transporter) interacting protein | Belardinelli et. al., 2019 |
| Rv3680 | N378D | | Increase (n = 1) No change (n = 1) | Increase (n = 2) | Gene involved in protecting *M. tuberculosis* from Glycerol and Nitric Oxide toxicity | Whitaker et. al., 2020 |
| Rv3758c | F209L | *proV* | Decrease | Increase | Role in the maintenance of osmoregulation within the phagosome | Gautam et. al., 2019 |
| Rv3901c | A38G | | Increase | No change | Role in virulence of *M. marinum* | Ruley et. al., 2004 |
| | T16I | | Increase | No change | | |

References for the supplementary table 3.

1. de Knegt GJ, Bruning O, ten Kate MT, et al. Rifampicin-induced transcriptome response in rifampicin-resistant Mycobacterium tuberculosis. Tuberculosis (Edinb) 2013; 93(1): 96-101.

2. Dhar N, McKinney JD. Mycobacterium tuberculosis persistence mutants identified by screening in isoniazid-treated mice. Proc Natl Acad Sci U S A 2010; 107(27): 12275-80.

3. Keren I, Minami S, Rubin E, Lewis K. Characterization and transcriptome analysis of Mycobacterium tuberculosis persisters. mBio 2011; 2(3): e00100-11.

4. Schuessler DL, Cortes T, Fivian-Hughes AS, et al. Induced ectopic expression of HigB toxin in Mycobacterium tuberculosis results in growth inhibition, reduced abundance of a subset of mRNAs and cleavage of tmRNA. Mol Microbiol 2013; 90(1): 195-207.

5. Wang R, Kreutzfeldt K, Botella H, Vaubourgeix J, Schnappinger D, Ehrt S. Persistent Mycobacterium tuberculosis infection in mice requires PerM for successful cell division. Elife 2019; 8.

6. Williams EP, Lee JH, Bishai WR, Colantuoni C, Karakousis PC. Mycobacterium tuberculosis SigF regulates genes encoding cell wall-associated proteins and directly regulates the transcriptional regulatory gene phoY1. J Bacteriol 2007; 189(11): 4234-42.

7. Nimmo C, Brien K, Millard J, et al. Dynamics of within-host Mycobacterium tuberculosis diversity and heteroresistance during treatment. EBioMedicine 2020; 55: 102747.

8. Mittal P, Sinha R, Kumar A, et al. Focusing on DNA Repair and Damage Tolerance Mechanisms in Mycobacterium tuberculosis: An Emerging Therapeutic Theme. Curr Top Med Chem 2020; 20(5): 390-408.

9. Lam JT, Yuen KY, Ho PL, et al. Truncated Rv2820c enhances mycobacterial virulence ex vivo and in vivo. Microb Pathog 2011; 50(6): 331-5.

10. Lagutkin D, Panova A, Vinokurov A, Gracheva A, Samoilova A, Vasilyeva I. Genome-Wide Study of Drug Resistant Mycobacterium tuberculosis and Its Intra-Host Evolution during Treatment. Microorganisms 2022; 10(7).

11. Pulido PA, Novoa-Aponte L, Villamil N, Soto CY. The DosR dormancy regulator of Mycobacterium tuberculosis stimulates the Na(+)/K (+) and Ca (2+) ATPase activities in plasma membrane vesicles of mycobacteria. Curr Microbiol 2014; 69(5): 604-10.

12. Baddam R, Kumar N, Wieler LH, et al. Analysis of mutations in pncA reveals non-overlapping patterns among various lineages of Mycobacterium tuberculosis. Sci Rep 2018; 8(1): 4628.

13. Williams KJ, Jenkins VA, Barton GR, Bryant WA, Krishnan N, Robertson BD. Deciphering the metabolic response of Mycobacterium tuberculosis to nitrogen stress. Mol Microbiol 2015; 97(6): 1142-57.

14. Shockey AC, Dabney J, Pepperell CS. Effects of Host, Sample, and in vitro Culture on Genomic Diversity of Pathogenic Mycobacteria. Front Genet 2019; 10: 477.

15. Sarkar R, Mdladla C, Macingwana L, et al. Proteomic analysis reveals that sulfamethoxazole induces oxidative stress in M. tuberculosis. Tuberculosis (Edinb) 2018; 111: 78-85.

16. Betts JC, Lukey PT, Robb LC, McAdam RA, Duncan K. Evaluation of a nutrient starvation model of Mycobacterium tuberculosis persistence by gene and protein expression profiling. Mol Microbiol 2002; 43(3): 717-31.

17. Xu W, DeJesus MA, Rucker N, et al. Chemical Genetic Interaction Profiling Reveals Determinants of Intrinsic Antibiotic Resistance in Mycobacterium tuberculosis. Antimicrob Agents Chemother 2017; 61(12).

18. Fang H, Yu D, Hong Y, Zhou X, Li C, Sun B. The LuxR family regulator Rv0195 modulates Mycobacterium tuberculosis dormancy and virulence. Tuberculosis (Edinb) 2013; 93(4): 425-31.

19. Calamita H, Ko C, Tyagi S, Yoshimatsu T, Morrison NE, Bishai WR. The Mycobacterium tuberculosis SigD sigma factor controls the expression of ribosome-associated gene products in stationary phase and is required for full virulence. Cell Microbiol 2005; 7(2): 233-44.

20. Fleischmann RD, Alland D, Eisen JA, et al. Whole-genome comparison of Mycobacterium tuberculosis clinical and laboratory strains. J Bacteriol 2002; 184(19): 5479-90.

21. Ahidjo BA, Kuhnert D, McKenzie JL, et al. VapC toxins from Mycobacterium tuberculosis are ribonucleases that differentially inhibit growth and are neutralized by cognate VapB antitoxins. PLoS One 2011; 6(6): e21738.

22. Colangeli R, Arcus VL, Cursons RT, et al. Whole genome sequencing of Mycobacterium tuberculosis reveals slow growth and low mutation rates during latent infections in humans. PLoS One 2014; 9(3): e91024.

23. Li W, Fan X, Long Q, Xie L, Xie J. Mycobacterium tuberculosis effectors involved in host-pathogen interaction revealed by a multiple scales integrative pipeline. Infect Genet Evol 2015; 32: 1-11.

24. Rachman H, Strong M, Ulrichs T, et al. Unique transcriptome signature of Mycobacterium tuberculosis in pulmonary tuberculosis. Infect Immun 2006; 74(2): 1233-42.

25. Fisher MA, Plikaytis BB, Shinnick TM. Microarray analysis of the Mycobacterium tuberculosis transcriptional response to the acidic conditions found in phagosomes. J Bacteriol 2002; 184(14): 4025-32.

26. Hampshire T, Soneji S, Bacon J, et al. Stationary phase gene expression of Mycobacterium tuberculosis following a progressive nutrient depletion: a model for persistent organisms? Tuberculosis (Edinb) 2004; 84(3-4): 228-38.
27. Belardinelli JM, Stevens CM, Li W, et al. The MmpL3 interactome reveals a complex crosstalk between cell envelope biosynthesis and cell elongation and division in mycobacteria. Sci Rep 2019; 9(1): 10728.

28. Whitaker M, Ruecker N, Hartman T, et al. Two interacting ATPases protect Mycobacterium tuberculosis from glycerol and nitric oxide toxicity. J Bacteriol 2020; 202(16).

29. Gautam US, Mehra S, Kumari P, et al. Mycobacterium tuberculosis sensor kinase DosS modulates the autophagosome in a DosR-independent manner. Commun Biol 2019; 2: 349.

30. Ruley KM, Ansede JH, Pritchett CL, Talaat AM, Reimschuessel R, Trucksis M. Identification of Mycobacterium marinum virulence genes using signature-tagged mutagenesis and the goldfish model of mycobacterial pathogenesis. FEMS Microbiol Lett 2004; 232(1): 75-81.

31. Catherine Vilchèze Mycobacterial Cell Wall: A Source of Successful Targets for Old and New Drugs http://dx.doi.org/10.3390/app10072278
